# QRAFT: Scaling coalescent-based species tree inference to one million taxa

**DOI:** 10.64898/2026.11.05.764586

**Authors:** Ashrafur Rahman, Anik Saha, Md. Shamsuzzoha Bayzid

**Affiliations:** Department of Computer Science and Engineering, Bangladesh University of Engineering and Technology, Dhaka-1205, Bangladesh

**Keywords:** Species tree estimation, Quartets, Multispecies coalescent, Consensus trees, Divide and conquer

## Abstract

Species tree inference from multi-locus data under the multispecies coalescent has not kept pace with phylogenomic datasets that now span hundreds of thousands of species. We present QRAFT, a statistically consistent method for estimating species trees by maximizing quartet support on phylogenomic datasets at unprecedented scales. For *n* taxa and *k* gene trees, QRAFT achieves near-linear running time of *O*(*kn* log^2^ *n*) when its recursion is balanced, which is within a polylogarithmic factor of the *O*(*kn*) time required simply to read the input. It also achieves asymptotically optimal *O*(*kn*) memory, so its memory requirements grow only linearly with the input size. QRAFT infers a million-taxon species tree from 1,000 genes in just three hours using only 298 GB of RAM, 64 CPU threads, and no GPUs. We estimate that the same analysis with ASTRAL-MP, a scalable parallel implementation of the widely used method ASTRAL, would take about 36 years (> 100, 000 times longer) and require prohibitive amounts of memory, far beyond practical computational limits. QRAFT maintains accuracy comparable to, and in many cases better than, leading species tree estimation methods. Notably, QRAFT reconstructed a microbial tree of life comprising 199,330 genomes in just a few hours using modest CPU resources. Thus, QRAFT brings highly accurate statistically consistent coalescent-based species tree inference to the unprecedented million-taxon regime and enables phylogenomic analyses towards the scale of the tree of life. QRAFT is publicly available at https://github.com/risenfromashes/qraft.

## 1 Introduction

International biodiversity initiatives are rapidly expanding genomic resources across hundreds of thousands of species, making phylogenomic analyses possible at unprecedented taxonomic scales. However, genome-scale phylogenomic analyses often reveal substantial discordance among gene trees which complicates the reconstruction of a species tree from multiple loci [1, 2]. Incomplete lineage sorting (ILS) is a major source of such discordance and is naturally modeled by the multispecies coalescent (MSC) [3–6].

Over the past decade, several summary methods have been developed for species tree inference from collections of gene trees, including methods that are statistically consistent under the MSC [7–16]. Among these, quartet-based methods such as ASTRAL and its variants have become widely used owing to their accuracy and favorable statistical properties [8, 17–19]. ASTRAL-III runs in *O*((*nk*)^1.726^*D*) time for *n* taxa and *k* gene trees, where *D* = *O*(*nk*) [20], and even with parallelism and randomization, ASTRAL-MP [21] remains practical only for datasets far smaller than hundreds of thousands of taxa. Quartet amalgamation methods (e.g., QFM and QMC families of methods [22–25]) recursively partition taxa to maximize quartet support. Their recent variants, TREE-QMC [26] and wQFM-TREE [27] work directly on gene trees rather than enumerating all Θ(*n*^4^) quartets, but still scale cubically in *n*. A different strategy, uDANCE [28] enables large-scale species tree inference by incrementally extending a backbone tree rather than inferring one *de novo*, but building the backbone remains difficult at large scales, and the method is not statistically consistent under the MSC.

Recent advances have challenged these longstanding scalability barriers of coalescent-based phylogenomics and substantially extended this frontier, with STELAR-X [29] and ASTRAL-X [30] scaling statistically consistent triplet- and quartet-based species-tree inference to 300,000 taxa. Yet these advances still leave open the challenge of globally optimizing quartet/triplet scores for species-tree inference at even larger scales. Here we present QRAFT (**Q**uartet **R**egrafting for **A**ccurate and **F**ast Species **T**ree Estimation), a new quartet-based approach that analyzes one million taxa in 2–5 hours using 64 CPU threads and no GPU. Remarkably, its running time grows quasilinearly with the number of taxa under balanced recursion. QRAFT also has an asymptotically optimal memory complexity of *O*(*nk*), which lets QRAFT run with memory proportional to the size of the input gene trees. At the same time, QRAFT matches or surpasses the accuracy of leading species tree estimation methods, including ASTRAL, across a wide range of datasets. QRAFT achieves this scalability by separating and scaling two problems—how to divide the taxon set and how to reconnect the resulting subtrees—that are intertwined in existing divide-and-conquer approaches. We propose several novel algorithmic techniques and data structures. Together, these techniques yield a new statistically consistent quartet-based method with *O*(*kn* log^2^ *n*) running time when the recursion is balanced and *O*(*kn*^2^ log *n*) in the worst case under highly imbalanced recursion, with *O*(*kn*+*pn*) memory on *p* workers. Extensive simulations demonstrate that QRAFT achieves an unprecedented scalability while maintaining accuracy comparable to, and often exceeding, that of state-of-the-art methods. We further demonstrate its practical utility by reconstructing a microbial Tree of Life comprising 199,330 genomes from 380 marker genes *de novo*, without relying on the backbone-extension approach employed by uDANCE [28]. QRAFT therefore combines statistical consistency, state-of-the-art accuracy, unprecedented scalability, and makes accurate species tree inference practical at the scale required for emerging tree of life initiatives.

## 2 Methods

Let *X* be a set of *n* taxa and let *G* = {*g*_1_, *g*_2_, …, *g_k_*} be a collection of *k* unrooted gene trees, where each *g_i_* may contain a subset *X_i_* ⊆*X* of the taxa and may be unresolved. For four distinct taxa *a, b, c, d*, the three resolved quartet topologies are *ab*|*cd, ac*|*bd*, and *ad*|*bc*. We write *Q*(*t*) for the set of resolved quartets displayed by a tree *t*, and *T* |*_Y_* for the restriction of a tree *T* to the set *Y* of taxa.

QRAFT addresses the Maximum Quartet Support Species Tree (MQSST) problem [8]. Given *G*, the objective is to find an unrooted species tree *T* on *X* that maximizes its total quartet agreement with the input gene trees: score 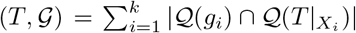. Thus, a gene-tree quartet contributes to the score exactly when the species tree displays the same topology on those four taxa. Under the multispecies coalescent model, maximizing this quartet-consistency objective yields a statistically consistent estimator of the species tree [8, 18].

### 2.1 Overview of QRAFT

QRAFT scales the MQSST objective to very large datasets by separating and scaling two decisions that divide-and-conquer quartet methods usually make together: how to divide the taxa, and how to reconnect the resulting subtrees. It proceeds in four stages (Fig. 1).

**Figure 1.**
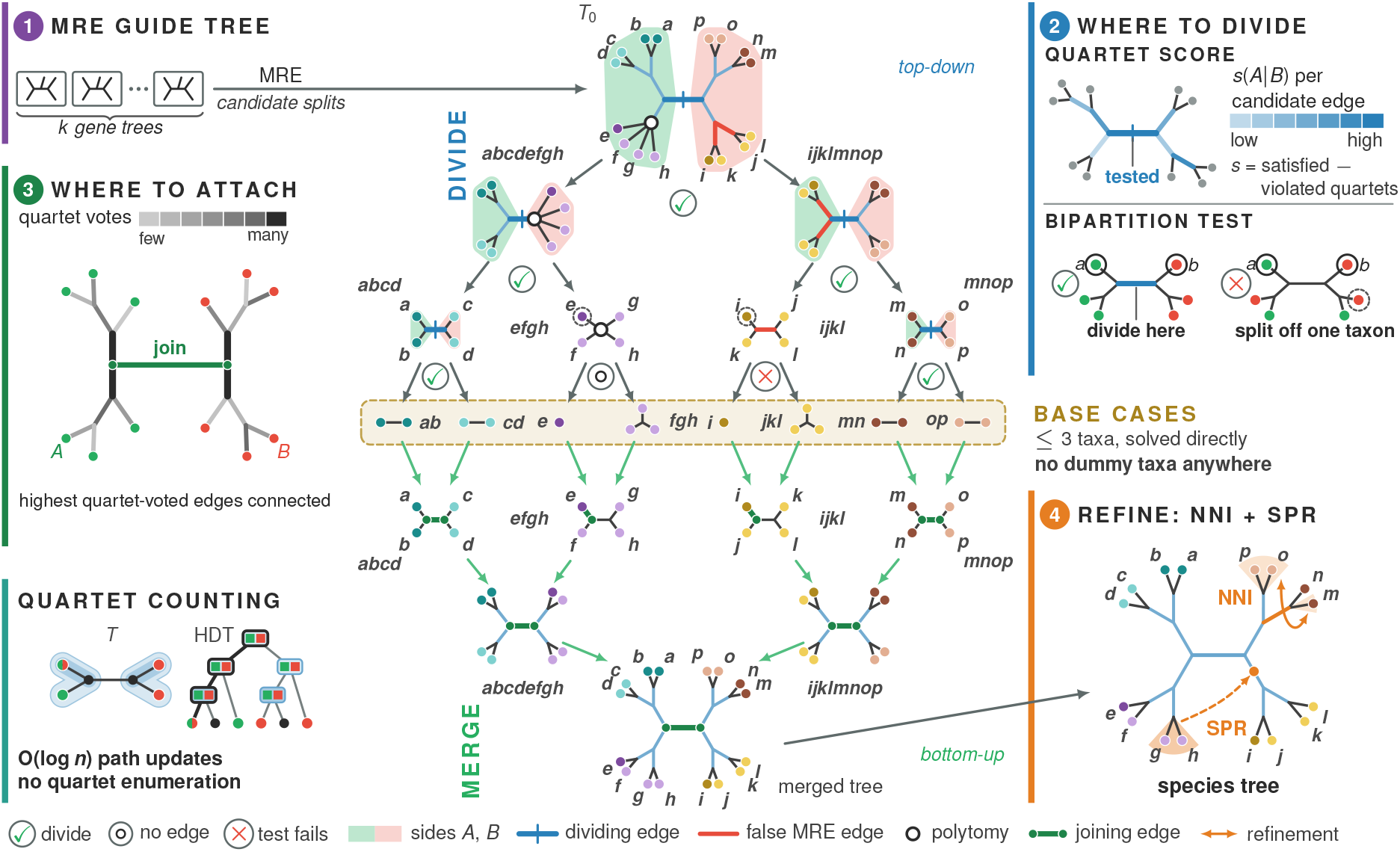
Overview of QRAFT. (1) MRE guide tree *T*_0_. (2) Top-down divide at the highest-scoring edge if it passes the bipartition test. (3) Bottom-up merge by regrafting at quartet-voted edges, with no dummy taxa. (4) NNI and SPR refinement.

1. **MRE guide tree.** QRAFT first builds an MRE consensus tree whose bipartitions serve as candidates for the search. Combining Wu’s algorithm [31] and the cluster identifiers of Jansson et al. [32], QRAFT constructs the MRE tree efficiently in optimal *O*(*kn*) space (Sect. 2.2).
2. **Quartet support guided recursive divide.** Guided by the MRE tree, QRAFT recursively selects bipartitions based on the “satisfied-minus-violated” quartet score. Since candidate splits from the MRE tree need not be a true species-tree edge, a bipartition test based on gene-tree internode distances is used to validate them before selection (Sect. 2.3).
3. **Contextual regrafting.** Unlike the traditional divide-and-conquer method such as QFM and QMC families that introduce artificial or dummy taxa, the divide step in QRAFT decides which taxa lie on each side of a bipartition, but not where the two subtrees join. QRAFT chooses the attachment edges only after both subtrees are built, using a novel quartet-based contextual regrafting technique, so no dummy or artificial taxa are needed. Importantly, we show that the optimal attachment edge on each side can be determined independently under the quartet objective, avoiding a joint search over pairs of edges. Subtrees are joined bottom-up until the full tree is assembled (Sect. 2.5).
4. **Refinement.** Finally, QRAFT uses bounded NNI and SPR moves to correct local errors (Sect. 2.6). Stages 2–4 all require many closely related quartet-score queries, and computing these scores by explicitly enumerating quartets would take Θ(*km*^4^) time per query on *m* taxa. QRAFT instead encodes each query as a coloring of the taxa and updates quartet counts incrementally on hierarchical decomposition trees (HDTs), using a small set of counters instead of answering each query from scratch. A whole family of queries is thus answered exactly in *O*(*km* log *m*) time (Sect. 2.7).

### 2.2 The MRE Starting Tree

The MRE consensus considers gene-tree clusters in decreasing order of frequency. It adds a cluster to the growing consensus tree whenever it is compatible with those already accepted [33].

To construct the MRE consensus tree *T*_0_ efficiently, we build on the algorithm of Wu [31]. Wu computes exact cluster identifiers with the dynamic set-equality structure of Gawrychowski et al. [34] in quasilinear time and space. We instead compute these identifiers with the divide-and-conquer labeling of Jansson et al. [32], which takes *O*(*kn* log *n*) time [32] and *O*(*kn*) space. Using the identifiers and counting sort, cluster frequencies are found within the same bound. Supplementary Section S1.1.1 gives a worked example.

Wu’s algorithm then starts from an unresolved star tree and refines it by inserting compatible most frequent clusters one at a time. For a candidate cluster *c*, we first use a link-cut tree structure to find the lowest vertex *u* of the current consensus tree whose descendant taxa contain all taxa in *c*. We then use balanced (AVL) search trees to check whether *c* is exactly the union of a set of complete child subtrees of *u*. If this condition holds, the cluster is compatible with the current consensus tree and is accepted to refine the tree [34]. Both data structures support queries and updates in *O*(log *n*) time, as in Wu’s algorithm, and following his analysis this part takes *O*(*kn* log^2^ *n*) time and *O*(*kn*) space. Combining both parts, the MRE construction takes *O*(*kn* log^2^ *n*) time and *O*(*kn*) space. Supplementary Section S1.1.2 presents the algorithm.

### 2.3 Quartet-Guided Recursive Divide

We now consider recursively dividing the taxon set *X* into smaller subproblems. QRAFT uses the bipartitions obtained from the MRE consensus tree *T*_0_ to guide this step. Starting from *X*, we repeatedly choose an edge of the current restriction of *T*_0_ and recurse on the two sides of the corresponding bipartition. The key question is therefore which bipartition to choose when several candidates are available.

Consider a subproblem on a taxon set *S*, where QRAFT selects a bipartition *A*|*B*. Since subsequent recursion and contextual regrafting may preserve this split, early choices can affect large portions of the final tree. We therefore score the candidate bipartitions by quartet support and choose the highest-scoring one.

Let *q* = *ab*|*cd* be a resolved gene-tree quartet and consider a candidate bipartition *A*|*B*. We say that *q* satisfies *A*|*B* if *a, b* lie on one side and *c, d* on the other. And *q* violates *A*|*B* if two taxa lie on each side but the quartet pairs taxa across the bipartition. Quartets with three or four taxa on the same side are ignored. Following the satisfied-minus-violated criterion of wQFM [35] and wQFM-TREE [27], we define *s*(*A*|*B*) = #{satisfying quartets across all gene trees} − #{violating quartets across all gene trees}. Thus, a larger value of *s*(*A*|*B*) indicates stronger quartet support for the candidate bipartition.

Accordingly, for a subproblem on taxon set *S*, QRAFT scores every nontrivial bipartition induced by the edges of *T*_0_|*S*. Explicitly enumerating quartets, however, would be prohibitively expensive. Instead, we express each score as a counting query. We color the two sides of the bipartition 0 and 1. A satisfying quartet then has pattern 00|11, whereas a violating quartet has pattern 01|01. Thus, writing *W* (*·*) for the corresponding gene-tree quartet count, *s*(*A*|*B*) = *W* (00|11) −*W* (01|01). The HDT-based counting procedure of Sect. 2.7 scores all the candidate edges together in *O*(*k*|*S*| log |*S*|) time. We then select the highest-scoring candidate, breaking ties in favor of the more balanced one.

A high quartet score, however, does not guarantee that the selected bipartition is a true species-tree edge. We therefore perform a bipartition test (Sect. 2.4) to guard this decision and preserve statistical consistency. If the candidate passes, we accept it. Otherwise, we use a trivial bipartition that separates one taxon from the rest. This cannot introduce an incorrect internal bipartition while still allowing the recursion to proceed.

### 2.4 The Bipartition Test

Our proposed bipartition test uses the internode distances between gene-trees used in ASTRID [36] and NJst [37]. But unlike these methods [38], our test stays statistically consistent under a random model of missing taxa (Definition 2.4 in Section 2.8).

#### Subpath between anchors

Assume that gene *g* has present taxa *X_g_* ⊆*X* and observed tree *H_g_* = *G_g_*|*X_g_*. For a candidate *A*|*B* of subproblem *S*, with |*A*|, |*B*| ≥ 2, fix two taxa as anchors *a* ∈ *A* and *b* ∈ *B*. Consider the *a*–*b* path between the anchors in *H_g_*. Every taxon *c* of *H*(*_g_* connects to this path at a vertex), say *u* (Fig. 2a). We define *h_g_*(*c*) as the length of the *a*–*u* subpath, 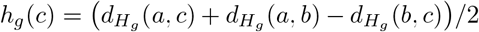. For distinct non-anchor taxa *x, y*, the shared genes are *G_xy_* = {*g* : {*a, b, x, y*} ⊆*X_g_*}, with count *k_xy_* = |*G_xy_*| > 0, we define the mean gap 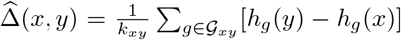. The mean gap is the central device of our test. It tells us about the species tree topology.

**Figure 2.**
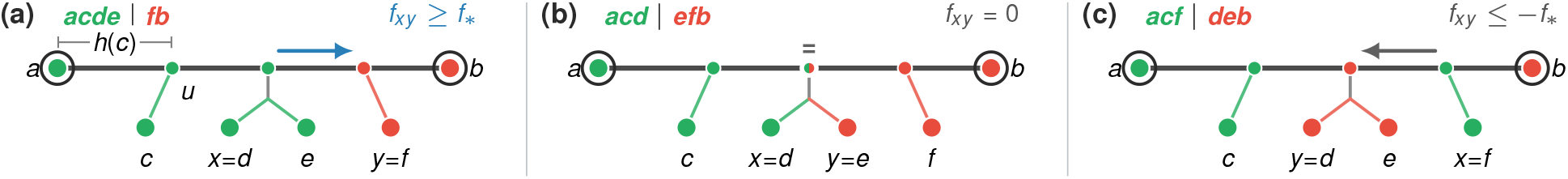
Limits of the mean gap, 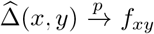. In each panel, *x* and *y* are the extreme pair *x*^*^ and *y*^*^ of the test, the last taxon of *A* and the first taxon of *B* on the *a*–*b* path. (a) True split: *A* attaches before *B* on the *a*–*b* path, *f_xy_* ≥*f*_*_. Taxon *c* attaches to the path at *u*, and *h*(*c*) is the length of the *a*–*u* subpath. (b) *x* and *y* branch off one vertex, *f_xy_* = 0. (c) Overlap, *f_xy_* ≤ −*f*_*_.

##### Lemma 2.1

(Mean gap sign characterization). *Given gene trees sampled under the model conditions* (*Definition 2.4), for distinct taxa a, b, x, y, there is a constant f*_*_ > 0 *such that* 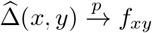*, that is, it converges in probability to f_xy_ as k → ∞, where f_xy_* ≥*f*_*_ *if the species quartet is ax*|*by, f_xy_* = 0 *if it is ab*|*xy, and f_xy_* ≤ −*f*_*_ *if it is ay*|*bx.*

On the species tree, the sign of the mean gap 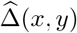 tells us the species tree quartet: positive mean gap implies the quartet *ax*|*by* (Fig. 2a), zero mean gap implies the quartet *ab*|*xy* (Fig. 2b) and negative mean gap implies the quartet *ay*|*bx* (Fig. 2c). For a false split there is always a quartet with *ab*|*xy* or *ay*|*bx*, and thus the mean gap cannot be strictly positive in the limit. This is the essence of our test.

#### The test

On the species tree, choose *x*^*^ ∈ *A \* {*a*} attaching last on the *a*–*b* path and *y*^*^ ∈ *B \* {*b*} attaching first (Fig. 2). Using a running search, QRAFT finds *x*^*^ by starting with any non-anchor taxon, say *x*_0_ ∈ *A \* {*a*}, and replacing it with *x* whenever 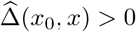. Similarly, *y*^*^ can be found in *B \* {*b*}. The test accepts *A*|*B* when 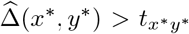, where the threshold 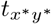 is chosen so that it tends to zero but more slowly than the sampling noise, such as 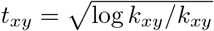.

If the *A*|*B* edge exists, the species quartet is *ax*^*^|*by*^*^, and therefore, by Lemma 2.1, with probability tending to one the mean gap is eventually strictly positive and greater than our threshold (Fig. 2a). For a false split, *x*^*^ and *y*^*^ either attach at the same vertex of the *a*–*b* path, inducing the quartet *ab*|*x*^*^*y*^*^ of case (b) (Fig. 2b), or *x*^*^ attaches later than *y*^*^, inducing the quartet *ay*^*^|*bx*^*^ of case (c) (Fig. 2c). Either way, the mean gap cannot be positive in the limit. See the details in Supplementary Lemma S3.6.

#### Running time

Finding *x*^*^ and *y*^*^ takes *O*(|*S*|) comparisons. Each comparison scans shared genes with indexed gene-tree distances, giving *O*(*k*|*S*|) total time using LCA queries without an *n* ×*n* distance matrix. Supplementary Section S3 gives the details.

### 2.5 Contextual Regrafting

The recursive divide gives rise to progressively smaller subproblems. Once the two child subproblems of a recursive call have been solved, it remains to determine how their resulting trees should be connected. Existing quartet-based divide-and-conquer methods often carry *dummy* or *artificial* taxa through the recursion to guide this merge step later on [25, 35, 39]. We instead observe that deferring the merge decision until both sides have been inferred allows the subproblems to remain free of dummy taxa. QRAFT therefore keeps the divide step simple and decides the attachment afterward by directly optimizing the quartet objective.

To determine this attachment, we introduce contextual regrafting. In essence, this can be viewed as a two-sided generalization of SPR [40] since both inferred trees are unrooted and the attachment edge is initially unknown on both sides. Apparently, finding the optimal merge requires a quadratic search over all pairs of attachment edges. However, we show that the quartet objective can be optimized separately on the two sides. As a result, the complete regrafting step can be carried out in *O*(*k*|*S*| log |*S*|) time.

#### The regrafting objective

Let us consider a subproblem on a taxon set *S* ⊆*X*, and suppose that the divide step chooses the bipartition *A*|*B*. Let *T_A_* and *T_B_* be the trees returned by the subproblems on *A* and *B*, respectively. To reconstruct a tree on *S*, we need to choose two edges *e_A_* in *T_A_* and *e_B_* in *T_B_*, subdivide them, and connect the newly formed vertices (Fig. 3a). We aim to choose these attachment edges so that the resulting tree has the highest quartet score on the gene trees restricted to *S*.

**Figure 3:**
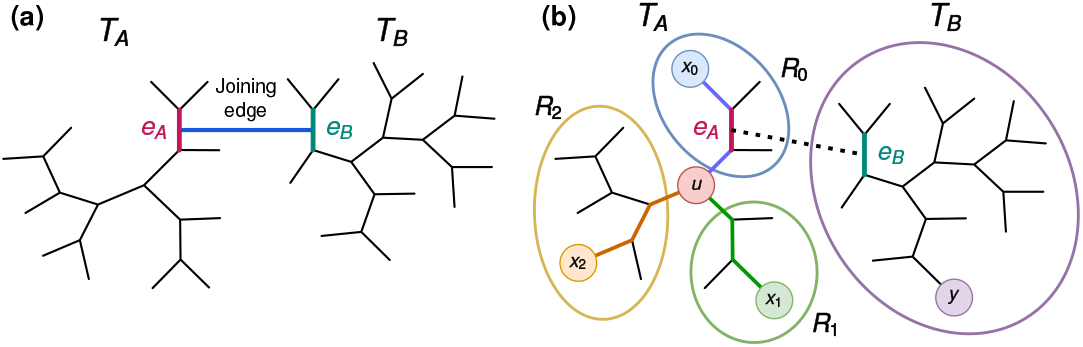
Contextual Regrafting. (a) Trees *T_A_* and *T_B_* are joined at independently chosen edges *e_A_* and *e_B_*. (b) Removing *u* yields regions *R*_0_, *R*_1_, *R*_2_. If *e_A_* is in *R*_0_, the quartet {*x*_0_, *x*_1_, *x*_2_, *y*} attains topology *x*_0_*y* |*x*_1_*x*_2_.

If both sides contain at least two taxa, there are (2|*A*|− 3)(2|*B*|− 3) possible pairs of attachment edges. A quadratic search over all pairs, however, would be too costly. Fortunately, we prove that under our quartet-score objective, the optimal edge for attaching *B* to *T_A_* can be determined without knowing where *A* will attach to *T_B_*, and vice versa (Lemma 2.2). Therefore, we can optimize the two sides independently instead of searching over all pairs of edges.

##### Lemma 2.2

(**Separability of the regrafting objective**). *The quartet score of the tree obtained by joining T_A_ and T_B_ at edges e_A_ and e_B_ can be written as c* + *ϕ_A_*(*e_A_*) + *ϕ_B_*(*e_B_*), *where c is independent of both attachment edges, ϕ_A_ depends only on e_A_, and ϕ_B_ depends only on e_B_. Consequently, the two attachment edges can be optimized independently.*

*Proof.* Consider any quartet of taxa from *S* = *A ∪ B*. Up to exchanging *A* and *B*, its taxa can be distributed across the two sides in only three ways: 4|0, 2|2, or 3|1.

**Case 1 (**4|0). All four taxa lie on the same side, say *A*. Their quartet topology is determined entirely by the corresponding subtree, here *T_A_*, and is therefore independent of both attachment edges.

**Case 2 (**2|2). Let *x*_0_, *x*_1_ ∈ *A* and *y*_0_, *y*_1_ ∈ *B*. Regardless of where the two trees are joined, the resulting tree displays *x*_0_*x*_1_ |*y*_0_*y*_1_, since the two taxa on each side meet within their respective subtree before reaching the joining edge. Hence, these quartets are again independent of *e_A_* and *e_B_*.

**Case 3 (**3|1). Let *x*_0_, *x*_1_, *x*_2_ ∈ *A* and *y* ∈ *B*. Since *y* is the only taxon in *B*, the quartet topology depends only on where *B* attaches to *T_A_*. The choice of *e_B_* cannot affect it. Thus, such quartets contribute only to a function *ϕ_A_*(*e_A_*). Symmetrically, quartets with three taxa in *B* contribute only to *ϕ_B_*(*e_B_*).

The first two cases contribute only a constant *c*, while the third contributes separately to *ϕ_A_*(*e_A_*) or *ϕ_B_*(*e_B_*). Therefore, the total quartet score has the form *c* + *ϕ_A_*(*e_A_*) + *ϕ_B_*(*e_B_*), which proves the claim. □

Thus we can optimize one side at a time. Even then, finding quartet-scores for every edge independently would be too costly. We now present how we evaluate all candidate edges together in *O*(*k*|*S*| log |*S*|) time.

#### Efficient optimization of a single side

We now focus on finding the best edge *e_A_* at which to attach *B* to *T_A_*. Optimizing *e_B_* in *T_B_* is symmetric. Consider an internal node *u* of *T_A_*, and let *R*_0_, *R*_1_, *R*_2_ be its three incident subtrees. Take taxa *x_r_* ∈ *R_r_* for *r* ∈ {0, 1, 2} and a taxon *y* ∈ *B* (Fig. 3b). Interestingly, the quartet induced by *x*_0_, *x*_1_, *x*_2_, *y* depends only on which *R_r_* contains the attachment edge *e_A_*. In particular, if *e_A_* lies in *R_r_*, then the quartet pairs *y* with *x_r_*, while the other two taxa form the opposite pair.

This allows us to obtain a local summary of the support for each attachment direction. For *r* ∈ {0, 1, 2}, let *q_r_*(*u*) denote the number of gene-tree quartets containing one taxon from each of *R*_0_, *R*_1_, *R*_2_ and one from *B* in which the taxon from *B* is paired with the taxon from *R_r_*. Thus, *q_r_*(*u*) measures the support at *u* for placing *e_A_* within *R_r_*. Using the four-color HDT queries of Sect. 2.7, all three values are obtained for every internal node in *O*(*k*|*A*| log |*A*|) time.

For an edge *e* of *T_A_*, let *r*(*u, e*) denote the branch of *u* reached by moving from *u* toward *e*. Its attachment score is 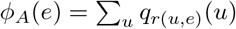. The key observation is that these scores need not be evaluated independently.

If adjacent edges *e* and *e*′ meet at *u*, directed toward *R_i_* and *R_j_*, respectively, then every internal node *v*≠ *u* sees both edges in the same branch. Hence, only the contribution at *u* changes, and *ϕ_A_*(*e*′) −*ϕ_A_*(*e*) = *q_j_*(*u*) −*q_i_*(*u*). Thus, after computing one edge score, a traversal of *T_A_* obtains all others in *O*(|*A*|) time. We choose the maximizing edge as *e_A_* and proceed symmetrically for *T_B_*, giving *O*(*k*|*S*| log |*S*|) time overall.

### 2.6 NNI and SPR Refinement

Contextual regrafting yields a tree with provable statistical consistency. However, the inference may leave local errors. We therefore apply NNI and SPR moves to improve the tree while retaining the guarantee. We now present an overview of these stages. Details are provided in Supplementary Section S1.2 and Figure S1.

#### NNI refinement

An NNI move compares the three possible arrangements around an internal edge. Existing approaches can require *O*(*kn*^2^) time [41]. Q-SPR [40] obtained the relevant quartet scores for one edge in *O*(*kn* log^2^ *n*) time but did not use them for NNI. QRAFT instead computes these scores for all internal edges together in *O*(*kn* log *n*) time using HDT-based counting (Sect. 2.7).

#### SPR refinement

SPR allows a subtree to move farther across the tree. For any fixed subtree, QRAFT can find its best destination exactly in *O*(*kn* log *n*) time using the same HDT framework. Repeating this for every subtree, however, would be quadratic. So we first screen promising subtrees using a small set of representative internal nodes. Only the top *c* candidates are then searched exactly over the full tree. With at most *z* representatives and *ρ* rounds, the complete SPR refinement thus takes *O*(*ρ*(*z* + *c*)*kn* log *n*).

### 2.7 HDT-based Quartet Counting

QRAFT repeatedly needs to solve the same problem: after assigning two or four colors to the taxa, counting gene-tree quartets of a few required colored topologies. Direct enumeration is prohibitively expensive, while ordinary DP on the gene trees does not support the many recolorings and progressively smaller subproblems that arise throughout QRAFT. We therefore build on the HDT framework of Brodal et al. [42–44].

#### Hierarchical decomposition trees (HDT)

An HDT reorganizes a gene tree into a locally balanced hierarchy of components. We use the four types of component of Brodal et al [43], each representing a set of nodes in the original tree. Since an HDT is locally balanced, changing the color of a leaf changes only logarithmically many components. Moreover, the Extract and Contractoperations of Brodal et al. [43] let us retain only the part relevant to the current active taxon set. Thus, both recoloring and subsequent queries scale with the current subproblem rather than with the full gene tree.

#### Counter states

At every HDT component, we store a constant number of counters (at most 23 for two colors and 56 for four colors) that summarize the colored configurations relevant to our quartet queries. QRAFT needs five quartet summaries: 00 | 11 and 01 | 01 for two colors, and 01 | 23, 02 | 13, and 03 | 12 for four colors. We therefore retain only the one-, two-, three-, and four-leaf configurations that can eventually contribute to these summaries. The complete counter definitions are given in Supplementary Table S1.

#### Traversal and updates

The counters define a DP over the HDT. They are computed by a bottom-up traversal of the HDT components using a fixed set of rules that obtain the counters of each component by combining or reusing those of its children in constant time. During a query-tree traversal (e.g., the consensus tree, or each side during regrafting), only changed colors are propagated and the remaining states reused, avoiding computation from scratch for every coloring, as in the Count routine of Brodal et al. [43].

Together, local balance, efficient recoloring, and contraction let one traversal answer a family of quartet queries in *O*(*km* log *m*) time for *m* active taxa. Further details are given in Supplementary Section S1.3.

### 2.8 Running Time, Memory, and Statistical Consistency

The following theorem gives the running time and memory of QRAFT (proof in Supplementary Section S1.4). **Theorem 2.3.** *For k single-copy gene trees and a recursion of height h, QRAFT runs in O*(*kn*(*h* + log *n*) log *n*) *time. Hence, the running time is O*(*kn* log^2^ *n*) *for a balanced recursion with h* = *O*(log *n*)*, and O*(*kn*^2^ log *n*) *in the worst case. With p workers, QRAFT uses O*(*kn* + *pn*) *memory.*

Let *X* be a fixed set of *n* ≥ 4 taxa, *X_g_* ⊆*X* the set of taxa present in gene *g*, and *G_g_* the corresponding gene tree. QRAFT observes *H_g_* = *G_g_*|*X_g_*.

#### Definition 2.4

(Model Conditions). We assume the following model conditions:

(M1) *Multispecies coalescent:* the gene trees *G_g_* are independent and identically distributed under the MSC on a binary species tree *σ* with positive internal branch lengths, and

(M2) *Full subset coverage:* the sets *X_g_* are independent and identically distributed and independent of the gene trees, with Pr(*B* ⊆*X_g_*) > 0 for every *B* ⊆*X* with |*B*| ≤ 4.

Condition (M1) is the multispecies coalescent model [45,46] on a species tree whose topology is identifiable from gene-tree frequencies [47], and (M2) is the full subset coverage model *M*_fsc_ of Nute et al. [48] with *k* = 4. Under these conditions, QRAFT is *statistically consistent* (Theorem 2.5; proof is in Supplementary S3).

#### Theorem 2.5

(Statistical consistency). *Under the model conditions* (*Definition 2.4*), *QRAFT is statistically consistent, for every starting tree, with or without NNI and SPR refinement*.

## 3 Results

We evaluated QRAFT on simulated and biological datasets and compared it with ASTRAL-IV, the ASTRAL implementation in ASTER [49], wQFM-TREE [27], ASTRAL-MP [21], STELAR-X 2.0.0 [29], and ASTRAL-X 1.0.0 [30]. The experiments ran on Linux servers with AMD EPYC processors, including the EPYC 7742 and EPYC 9354, and NVIDIA RTX 4090 GPUs for the GPU-based methods (Supplementary Section S2.1). Most analyses were allocated a maximum running time of 48 hours and 128 GB of RAM. The scalability analyses with up to one million taxa or gene trees were allocated a maximum running time of 24 hours and 328 GB of RAM.

### Simulated Datasets

For the scalability analyses, we generated gene trees under the multispecies coalescent (MSC) following the STELAR-X benchmark settings and ILS model conditions [29, 30] using Sim-Phy [50], which we substantially re-engineered and optimized to support simulations with up to one million taxa (SimPhy-X [51]). We varied the number of taxa from 1,000 to 1,000,000 with 1,000 gene trees and the number of gene trees over the same range with 1,000 taxa. All datasets were simulated under four levels of incomplete lineage sorting (ILS), from low (ILS-L1) to medium-high (ILS-L4). To further evaluate accuracy, we used the 37-taxon mammalian dataset [52, 53], 200-, 500- and 1,000-taxon datasets covering the eight model conditions of ASTRAL-II [17], and the 10,000-taxon ASTRAL-MP dataset [21] (details in Supplementary Section S2.4).

### Biological Datasets

We analyzed two biological datasets: the bacterial and archaeal dataset analyzed by uDANCE [28] comprising 199,330 taxa and 380 gene trees, the angiosperm dataset [54] with 9,524 taxa and 353 genes. Results on the angiosperm dataset are presented in (Supplementary Section S2.8.1).

We run QRAFT in the default refinement mode for scalability and accuracy, and the extended mode for biological datasets. Supplementary S2.3 and S2.9 give comparison between the modes and ablation studies.

### 3.1 Results on Scalability Analysis

We performed experiments on simulated datasets with large numbers of taxa and gene trees under medium ILS (ILS-L2) and reported running time, CPU memory usage, averaged over five replicates. In the first set of experiments, we fixed the number of gene trees at *k* = 1,000 and varied the number of taxa *n* from 1,000 to 10,000. Within our resource limits, ASTRAL-MP could be run only up to 10,000 taxa. On these datasets, QRAFT, ASTRAL-X, and STELAR-X were the fastest methods, completing the analyses in under a minute (Fig. 4a). Although ASTRAL-X and STELAR-X were slightly faster, they required 4.5–8.0× more CPU memory than QRAFT (Fig. 4a). At 10,000 taxa, QRAFT completed the analysis in 55 seconds using only 2.0 GB of RAM, compared with 16 and 15 seconds and 8.9 and 9.1 GB for ASTRAL-X and STELAR-X, respectively. ASTRAL-MP took 9.4 hours (613× and 2,200× the running times of QRAFT and STELAR-X, respectively) and used 126 GB of RAM (64× QRAFT’s memory usage). ASTER did not finish within 48 hours. Both ASTRAL-MP and ASTER exhibit more than quadratic growth in running time with increasing numbers of taxa as evident from Fig. 4a as well. At this relatively small scale of up to 10,000 taxa, constant-factor overheads dominated, and QRAFT’s asymptotic advantage in running time was not yet fully realized. In this regime, effective GPU parallelization gave STELAR-X and ASTRAL-X lower practical overheads than QRAFT. As dataset size increased, however, QRAFT became substantially faster than both methods from 75,000 taxa onward, consistent with our theoretical running time analyses (Figs. 4 and S4 in Supplementary Section S2.2).

**Figure 4.**
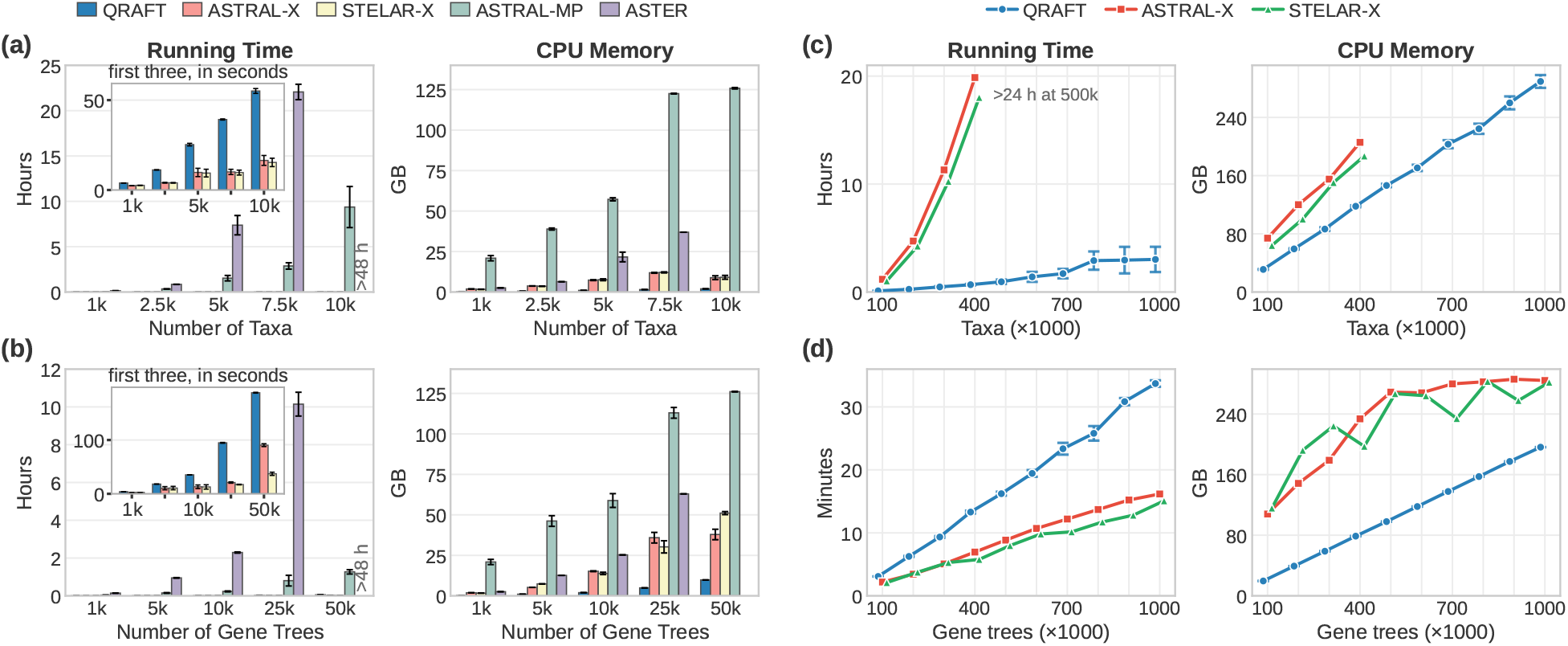
Running time and peak CPU memory. (a) 1,000 gene trees, 1,000–10,000 taxa. (b) 1,000 taxa, 1,000–50,000 gene trees. In (a,b), means over five replicates, ASTER exceeded 48 hours at 10,000 taxa and 50,000 gene trees, and the inset subfigures show seconds for QRAFT, STELAR-X and ASTRAL-X. (c) 1,000 gene trees, 100,000–1,000,000 taxa. (d) 1,000 taxa, 100,000–1,000,000 gene trees. In (c,d), ASTRAL-X and STELAR-X: replicate 1, default search space S1; QRAFT: means over three to five replicates per size.

Next, we evaluated scalability with respect to the number of genes while fixing the number of taxa at *n* = 1,000 (Fig. 4b). QRAFT was substantially faster than ASTRAL-MP and ASTER but was generally slower than STELAR-X and ASTRAL-X, while using substantially less memory than GPU-based methods (STELAR-X and ASTRAL-X). With 50,000 genes, ASTRAL-X and STELAR-X took 90 and 37 seconds and used 37.9 and 51.1 GB of RAM, respectively, whereas QRAFT took 3.1 minutes and used just 9.8 GB. ASTRAL-MP took 76 minutes and used 126 GB of RAM, and ASTER did not finish within 48 hours. Thus, STELAR-X, ASTRAL-X, and QRAFT all scaled well to large numbers of genes. As Figs. 4a,b show, their memory requirements grew linearly, consistent with their theoretical bounds.

### Scaling up the number of taxa and gene trees

To assess performance at larger scales, we evaluated QRAFT on datasets with up to one million taxa or genes. In the first experiment, we fixed *k* = 1,000 and increased *n* from 100,000 to 1,000,000 (Fig. 4c). These experiments ran on an AMD EPYC 9354, which was faster than the EPYC 7742, with 64 threads per run (Supplementary Section S2.1). STELAR-X and ASTRAL-X could not be run beyond 400,000 taxa within our resource budget.

QRAFT’s computational advantage over existing methods becomes increasingly pronounced beyond 100,000 taxa (Fig. 4c). At 400,000 taxa, ASTRAL-X and STELAR-X required 19.9 and 18.0 hours, respectively, whereas QRAFT completed the same analyses in only 50 minutes, a 22–24× speedup, while using approximately 40% less memory. Neither ASTRAL-X nor STELAR-X completed the 500,000-taxon analysis within 24 hours (Fig. 4c). In contrast, QRAFT scaled to one million taxa in an average of 3 hours, using at most 298 GB of RAM, a scale beyond the reach of existing statistically consistent summary methods.

As the number of taxa increased tenfold, from 100,000 to one million, QRAFT’s mean running time increased by only 27×, consistent with its theoretical subquadratic running time and well below the 100× increase expected under quadratic scaling (as clear from Fig. 4c). By comparison, the running times of ASTRAL-X and STELAR-X increased by approximately 17× between 100,000 and 400,000 taxa, indicating near-quadratic empirical growth (Fig. 4c). QRAFT’s empirical memory usage scaled nearly linearly (Figs. 4c,d), consistent with its theoretical memory complexity.

Extrapolating from the observed running time and memory growth, we estimated that ASTRAL-X and STELAR-X would require approximately five days and about 450 GB of RAM to analyze one million taxa. Our estimated computational demands of conventional quartet-based inference, specifically ASTRAL-MP, were considerably greater: approximately 36 years, with memory requirements exceeding practical hardware limits, making such an analysis infeasible even on high-performance computing infrastructure. With subquadratic running time scaling and linear memory growth, QRAFT reduced analyses estimated to require decades with conventional state-of-the-art methods such as ASTRAL to hours on a CPU-based system with moderate memory requirements, without requiring GPUs.

Next, we evaluated scalability with respect to the number of genes while fixing the number of taxa at *n* = 1,000 (Fig. 4d). All three methods scaled well to large numbers of genes, completing analyses of one million genes in 15–35 minutes. QRAFT was slower than STELAR-X and ASTRAL-X but used substantially less memory. It took 1.4–2.1× as long as ASTRAL-X and used about 31–82% less memory. With one million genes, QRAFT took 34 minutes and used 196 GB of RAM, ASTRAL-X took 16 minutes and used 284 GB, and STELAR-X took 15 minutes and used 282 GB. QRAFT thus scaled similarly to STELAR-X and ASTRAL-X with the number of genes, while using substantially less memory.

### 3.2 Results on Accuracy of Inference

We conducted experiments on a wide range of simulated datasets with varying model conditions. Remarkably, on the 37-taxon biologically-based mammalian simulated dataset, QRAFT achieved the best overall accuracy among all methods, including the state-of-the-art ASTRAL and wQFM-TREE, across all model conditions with varying levels of ILS and gene tree estimation error controlled by sequence lengths (Fig. 5a). QRAFT achieved the highest accuracy in three of the five model conditions: moderate (1X) and low (2X) ILS, and true gene trees. wQFM-TREE achieved the highest accuracy under high ILS (0.5X) and with 1,000-bp sequences. On the ASTRAL-II datasets (Fig. 5b), QRAFT and wQFM-TREE achieved the highest accuracy, significantly outperforming the other methods including ASTRAL-MP (*p* < 10^−9^).

**Figure 5.**
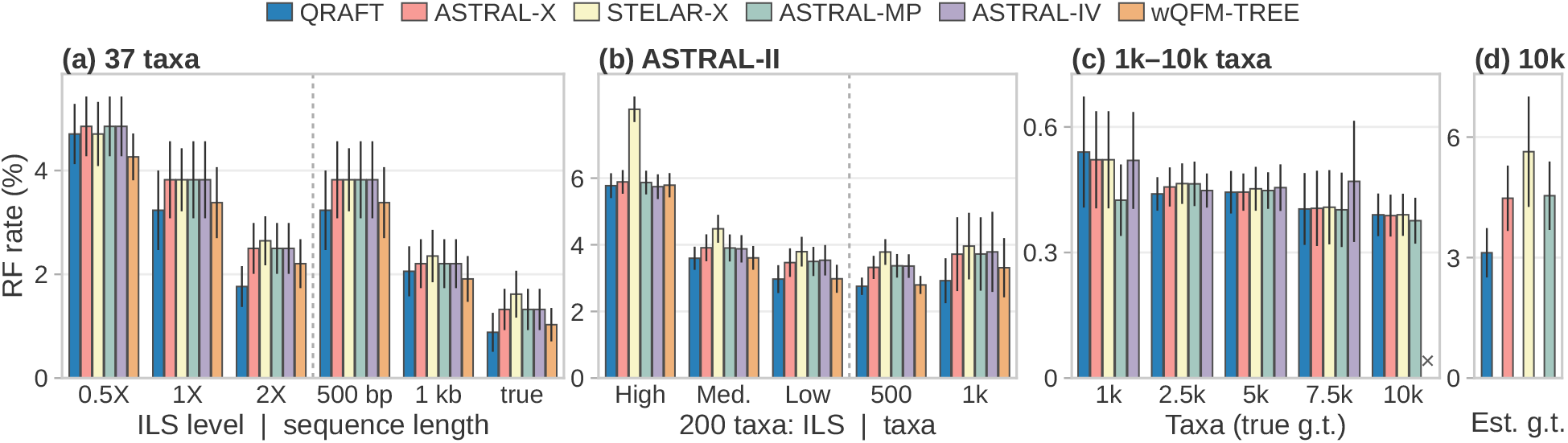
Mean RF rates. (a) 37 taxa across ILS levels and sequence lengths, and true gene trees. (b) 200 taxa at high, medium and low ILS, and 500 and 1,000 taxa. (c) 1,000–10,000 taxa, true gene trees (×: ASTRAL-IV exceeded 48 hours at 10000 taxa). (d) 10000 taxa, estimated gene trees.

To further assess accuracy at larger scales, we evaluated QRAFT on datasets comprising 1,000–10,000 taxa from the STELAR-X study (Fig. 5c). All methods achieved comparable accuracy, with no statistically significant differences. RF rates remained low on this dataset because we used true gene trees under moderate ILS. We also evaluated the 10,000-taxon dataset with estimated gene trees from the ASTRAL-MP study, where QRAFT achieved the highest accuracy, with an RF error rate of 3.1%, compared with 4.5% for both ASTRAL-MP and ASTRAL-X and 5.6% for STELAR-X (Fig. 5d). We further evaluated the accuracy on ultra-large dataset with up to one million taxa analyzed in Section 3.1 (see Supplementary Section S2.2). Together, these results demonstrate that QRAFT maintains state-of-the-art accuracy while achieving unprecedented scalability to the million-taxon regime.

### 3.3 Constructing the microbial tree of life

We re-analyzed the large-scale microbial dataset of Balaban et al. [28], comprising 199,330 genomes. Starting from the Web of Life (WoL) tree of 10,575 genomes, Balaban et al. [28] first incorporated additional genomes to obtain a backbone of 15,953 taxa (the 16K tree), which was subsequently expanded to the full dataset. In each expansion step, uDANCE placed new genomes onto the existing backbone using APPLES-2 [55], partitioned the resulting tree into smaller subsets (final 200K analysis used 78 partitions), inferred gene trees within each subset using RAxML-NG, and reconstructed local species trees using constrained ASTRAL before merging them into a single tree without ever globally optimizing the whole tree.

Unlike uDANCE, QRAFT reconstructs and optimizes the entire 199,330-taxon species tree using all 380 gene trees. Maximum-likelihood gene tree estimation with RAxML [56] is computationally prohibitive at this scale, so we inferred gene trees from the released marker alignments using FastTreeMP [57] under *LG* + Γ. We evaluated QRAFT in two modes: i) *de novo*, which infers the tree directly from the gene trees, and ii) guided *de novo*, which uses smaller trees only to construct a better starting tree. Specifically, the 16K tree estimated by QRAFT was used to refine the initial MRE tree on 200K taxa by excluding conflicting clusters and resolving polytomies. Subsequently, QRAFT performs global species tree optimization over all 199,330 taxa, without restricting the search to the guide topology. Remarkably, QRAFT completed the full analysis in just 5.55 hours de novo and 3.80 hours in guided de novo mode, including the intermediate analyses (Table S4). Including gene tree estimation with FastTreeMP (1,180 CPU hours), the complete QRAFT pipeline used about 1,535 CPU hours *de novo* and 1,423 guided, more than 200× less compute than the 334,344 CPU hours reported for uDANCE. This difference is particularly notable because the two methods solve problems of very different size: uDANCE estimates gene trees and species trees only within partitions of a few thousand genomes, whereas QRAFT optimizes the quartet score globally over all 199,330 genomes.

Fig. 6 shows the published tree beside QRAFT’s guided tree, and Fig. S6 in Supplementary S2.5 shows the de novo tree without guidance. Both QRAFT trees achieved higher quartet scores than the uDANCE tree, with gains of 9.49 × 10^18^ and 9.64 × 10^18^ for the de novo and guided analyses, respectively. QRAFT’s trees also agreed more closely with the GTDB taxonomy [58, 59]. Among the 153,559 genomes with a GTDB label, QRAFT recovered more monophyletic groups than uDANCE at every taxonomic rank (Table 1).

**Table 1.**
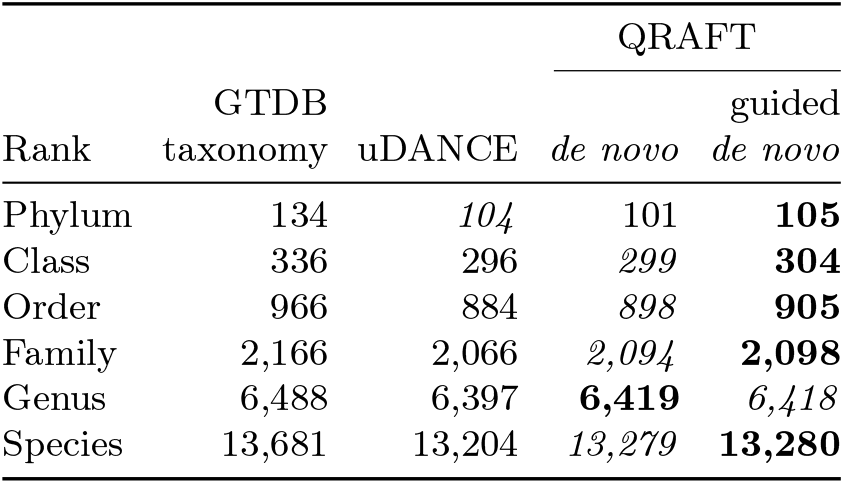
Monophyletic groups at every taxonomic rank according to GTDB taxonomy.

| Rank | GTDB taxonomy | uDANCE | QRAFT |  |
|---|---|---|---|---|
|  |  |  | *de novo* | guided *de novo* |
| Phylum | 134 | *104* | 101 | **105** |
| Class | 336 | 296 | *299* | **304** |
| Order | 966 | 884 | *898* | **905** |
| Family | 2,166 | 2,066 | *2,094* | **2,098** |
| Genus | 6,488 | 6,397 | **6,419** | *6,418* |
| Species | 13,681 | 13,204 | *13,279* | **13,280** |

**Figure 6.**
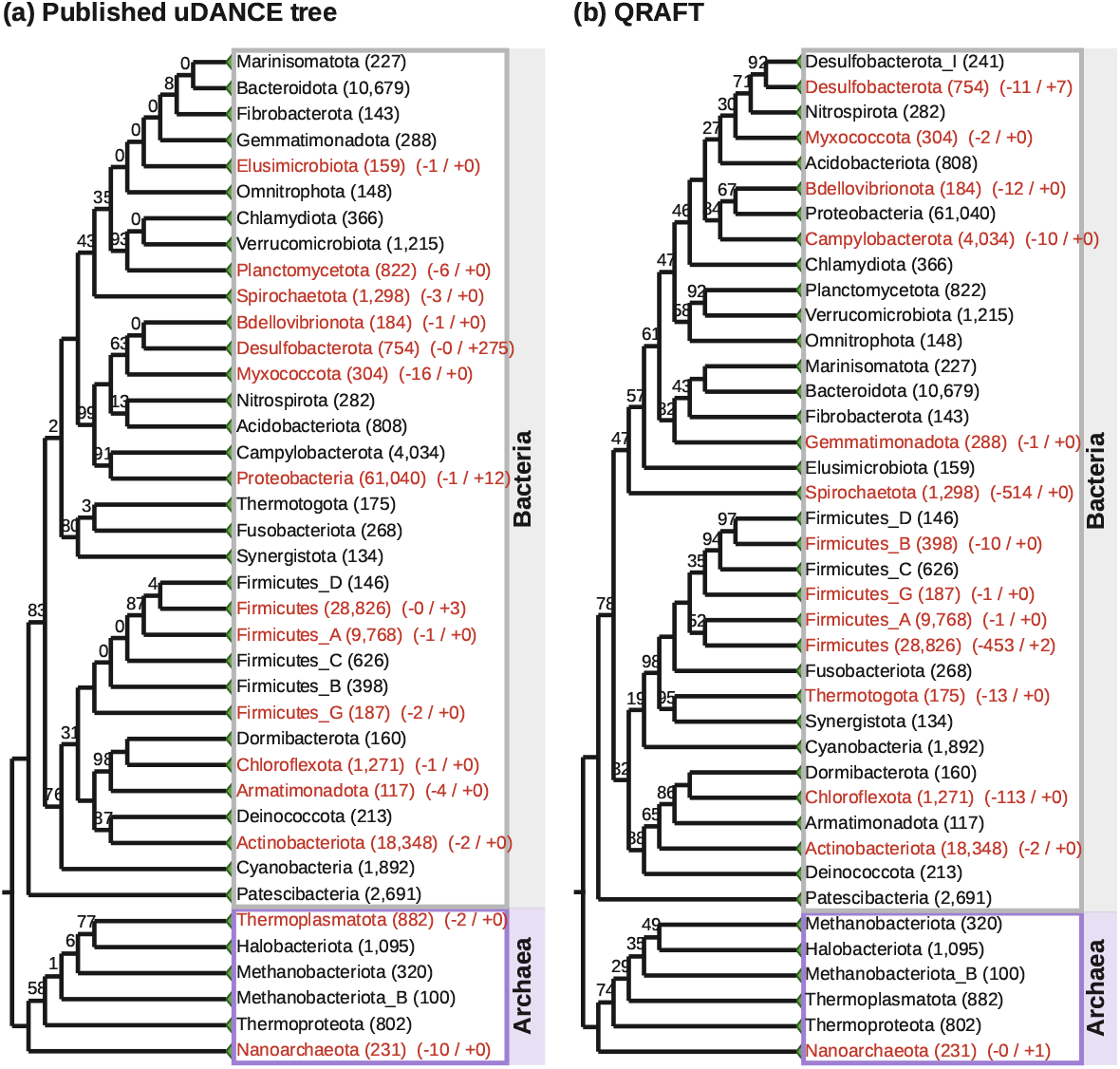
Microbial tree of life with 199,330 taxa. (a) Published tree [28]. (b) QRAFT’s guided *de novo* tree, rooted between Archaea and Bacteria, with the 40 GTDB phyla having at least 100 genomes collapsed. Local posterior supports [41] are 100% unless noted. Red labels: nonmonophyletic groups, with −k/+m members outside/intruders inside the best clade.

QRAFT recovered major evolutionary relationships consistent with previous studies, including the separation of Archaea and Bacteria, the basal placement of CPR within Bacteria, and the placement of DPANN as the earliestdiverging archaeal lineage (see Supplementary Section S2.6).

Importantly, these results were obtained with FastTree gene trees, which are generally less accurate than RAxML gene trees. In a separate analysis of the WoL1 dataset, QRAFT recovered 63 GTDB phyla from RAxML gene trees but only 60 from FastTree gene trees of the same markers (Supplementary Section S2.7).

## 4 Discussion

We introduced QRAFT, the first statistically consistent species tree inference method to reach the milliontaxon scale in just three hours using less than 300 GB of RAM and no GPUs. QRAFT introduces a new algorithmic framework that separates recursive partitioning from subtree reconnection, combining a novel quartet-based contextual regrafting technique and a new HDT-based quartet counting. Despite its unprecedented scalability, QRAFT maintains state-of-the-art accuracy and outperforms widely used methods such as ASTRAL under several challenging evolutionary conditions. We further reconstructed a 199,330-genome microbial Tree of Life de novo, representing, to our knowledge, the largest empirical phylogeny inferred using a statistically consistent method. Overall, QRAFT marks a major step towards accurate, statistically consistent phylogenomic inference at the scale of the Tree of Life.

## Supplementary material

## S1 Supplementary Methods

### S1.1 MRE Starting Tree

#### S1.1.1 Example: Exact Cluster Identifiers and Frequency Counting

We illustrate how QRAFT identifies equal clusters across different gene trees and obtains their frequencies without storing the clusters as explicit taxon sets. The identifiers are computed with the divide-and-conquer labeling of Jansson et al. [1] (their procedure Fast Label Trees). For simplicity, consider three gene trees on the taxa

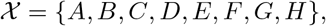

with integer identifiers 0, …, 7 assigned in this order. After applying the fixed rooting used by the MRE construction, suppose the trees are

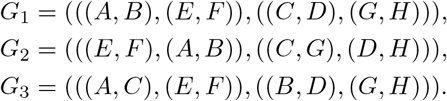

Consider the clusters induced by the left child of the root:

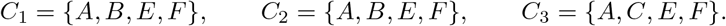

Thus, *C*_1_ and *C*_2_ represent the same cluster although they arise from different gene trees, whereas *C*_3_ is different. The goal is to discover this equality using compact integer identifiers.

The identifiers are constructed recursively by dividing the taxon range into halves. In the implementation, a range containing at most 64 taxa is represented directly by a 64-bit mask. To expose the recursion in this small example, we instead continue until each range contains at most two taxa.

##### First recursion level

The full taxon range is

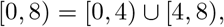

corresponding to

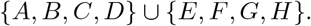

Each cluster is therefore represented by its intersection with these two halves:

| Cluster occurrence | [0, 4) | [4, 8) |
|---|---|---|
| *C*<sub>1</sub> | {*A*, *B*} | {*E*, *F*} |
| *C*<sub>2</sub> | {*A*, *B*} | {*E*, *F*} |
| *C*<sub>3</sub> | {*A*, *C*} | {*E*, *F*} |

At this stage, the two pieces are not yet stored as taxon sets. Instead, recursive calls assign exact identifiers to the distinct pieces occurring in each half.

##### Identifiers in the left half

The range [0, 4) is divided again as

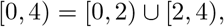

corresponding to {*A, B*} and {*C, D*}. The two distinct pieces that occur above decompose as

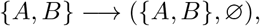

and

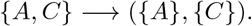

At the base range [0, 2), the occurring subsets are {*A*} and {*A, B*}. Using the local bit-mask representation

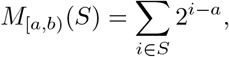

their masks are

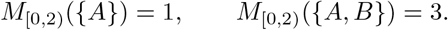

Ranking the distinct masks gives, for example,

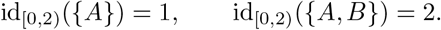

Similarly, for [2, 4) we may assign

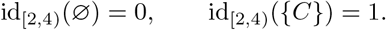

Returning to [0, 4), the two relevant subsets are now represented entirely by pairs of child identifiers:

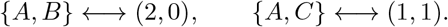

After sorting these pairs, each distinct pair receives a new identifier. For example,

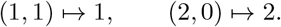

Hence,

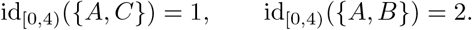

##### Identifiers in the right half

All three cluster occurrences have the same right piece, {*E, F*}. Dividing

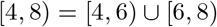

represents this subset as

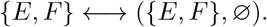

After assigning base identifiers and ranking the resulting pair, let

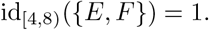

##### Final identifiers

We now return to the original range [0, 8). The three cluster occurrences are represented by

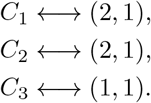

Sorting these pairs places the two equal representations together:

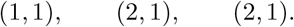

Assigning a new identifier to each distinct pair may therefore give

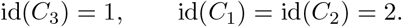

The numerical values of the identifiers themselves are immaterial. Their purpose is only to satisfy

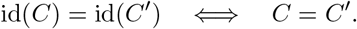

##### Frequency counting

Once the exact identifiers are available, cluster frequencies can be obtained without inspecting the taxon sets again. Each cluster occurrence is stored as a record containing its size, identifier, source gene, and source edge. For the three occurrences above, the relevant part of the records is

| Gene | Cluster size | Identifier | Cluster |
|---|---|---|---|
| *G*<sub>1</sub> | 4 | 2 | {*A*, *B*, *E*, *F*} |
| *G*<sub>2</sub> | 4 | 2 | {*A*, *B*, *E*, *F*} |
| *G*<sub>3</sub> | 4 | 1 | {*A*, *C*, *E*, *F*} |

A radix sort groups equal identifiers together. A single scan then counts the number of distinct gene trees contributing each identifier:

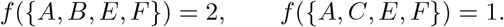

Thus, the first cluster is recognized as the same taxon set in *G*_1_ and *G*_2_ and receives frequency two, whereas the second occurs only in *G*_3_. The same procedure is applied simultaneously to all cluster occurrences extracted from the input gene trees. A final radix sort orders the distinct clusters by decreasing frequency for the subsequent MRE compatibility scan.

Importantly, no dictionary over arbitrary subsets of *X* is maintained. At each recursion level, only the cluster pieces that actually occur in the gene trees are represented, and their identifiers are obtained by sorting pairs of identifiers returned by the two recursive halves. Since the total number of such records at any level is *O*(*kn*) and there are *O*(log *n*) levels, the exact identifiers are obtained in *O*(*kn* log *n*) time using *O*(*kn*) space.

#### S1.1.2 Construction of the MRE Starting Tree

We present the construction of the MRE consensus tree in Algorithm 1. The accompanying divide-and-conquer procedure used to assign exact identifiers to gene-tree clusters is given in Algorithm 2.

### S1.2 NNI and SPR Refinement

After the recursive divide and contextual regrafting, QRAFT has reconstructed a complete binary tree. We next apply NNI and SPR moves as a post-processing step to correct remaining local errors. NNI changes only a small neighborhood of the current tree while SPR moves can relocate a subtree much farther in the tree. In the default mode, QRAFT applies a small number of carefully gated NNI moves. In the extended mode, we apply batched NNI and SPR moves iteratively.

#### S1.2.1 NNI refinement

Let us consider an internal edge *pq* of the current tree. Let *W* and *X* be the two subtrees incident to *p*, and let *Y* and *Z* be the two subtrees incident to *q* (Fig. S1a). Thus, the current arrangement around the edge is *WX* |*Y Z*. There are two alternative NNI arrangements, *WY* |*XZ* and *WZ* |*XY*, obtained by exchanging one of the neighboring subtrees across the edge. Thus, at every internal edge, there are three local arrangements to compare.

Interestingly, most quartets in the tree are unaffected by an NNI move. A quartet can change its topology only if it contains one taxon from each of the four subtrees, *w* ∈ *W*, *x* ∈ *X, y* ∈ *Y*, *z* ∈ *Z*. For these quartets, the current arrangement displays *wx*|*yz*, whereas the two NNI alternatives display *wy*|*xz* and *wz*|*xy*. Therefore, the gain of an NNI move can be obtained simply by comparing the gene-tree support for these three quartet topologies.

We compute these supports using a four-color query. Assigning colors 0, 1, 2, 3 to *W*, *X, Y*, *Z*, respectively, gives the three counts

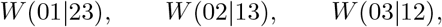

corresponding to *WX*|*Y Z, WY* |*XZ, WZ*|*XY*. For example, the gain from replacing *WX*|*Y Z* by *WY* |*XZ* is *W* (02|13) − *W* (01|23).

##### Algorithm 1

Construction of the MRE Starting Tree

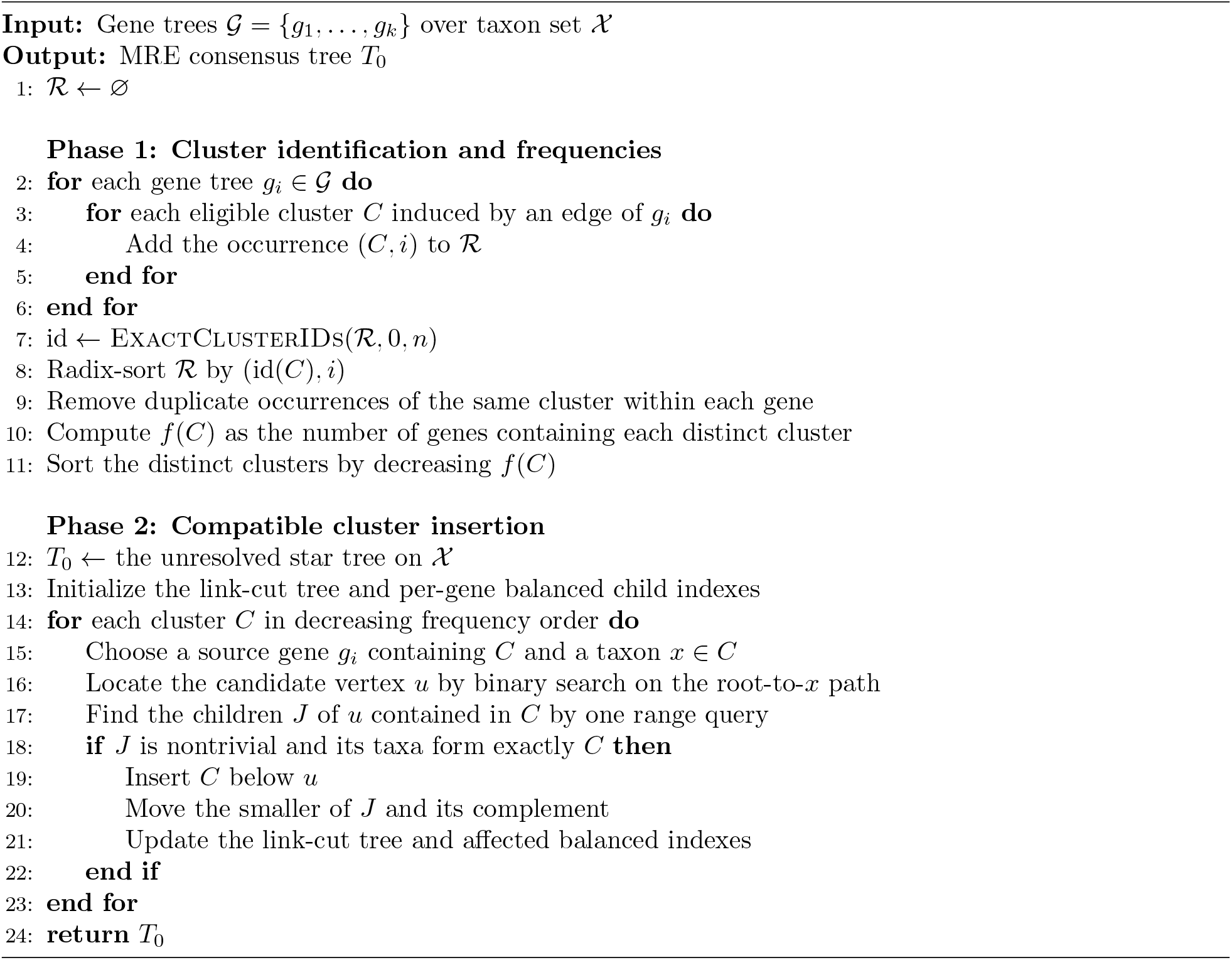

Importantly, these counts are not computed independently for every internal edge. As described in Sect. S1.3, QRAFT traverses the current tree while updating the corresponding four-color queries, thereby obtaining the three quartet counts for all internal edges together in *O*(*kn* log *n*) time. We then examine the two alternatives at every edge and select the NNI move with the largest quartet-score gain.

If the selected move is accepted, QRAFT applies the corresponding local exchange and continues. Fortunately, an NNI move affects only a small neighborhood of the modified edge. For any sufficiently distant edge, the same four surrounding subtrees remain in place, and therefore its previously computed NNI score is unchanged. Consequently, we need to recompute scores only for edges within distance two of the applied move. After these local updates, QRAFT again selects the move with the largest gain and repeats the process. Since the number of accepted moves is bounded by a constant and only a local neighborhood is rescored after each move, the complete NNI refinement remains within *O*(*kn* log *n*) time.

QRAFT can also apply several compatible NNI moves in the same refinement step. Once the gains of all candidate moves have been computed, we identify a set of positive-gain moves that can be performed simultaneously without interfering with one another. In particular, moves whose central edges share a node are not selected together, and nearby moves are combined only when the edge separating them remains unchanged. Under these conditions, the individual quartet-score gains can be added. We therefore use a dynamic program on the tree to select the compatible set with the largest total gain and apply all of its moves together. This allows several independent local errors to be corrected from a single *O*(*kn* log *n*) scoring traversal.

Every move is applied only if its exact quartet-score gain is positive, so refinement keeps the species tree once reached (Lemma S3.8.1, Sect. S3).

##### Algorithm 2

Exact Cluster Identifiers by Divide-and-Conquer Labeling

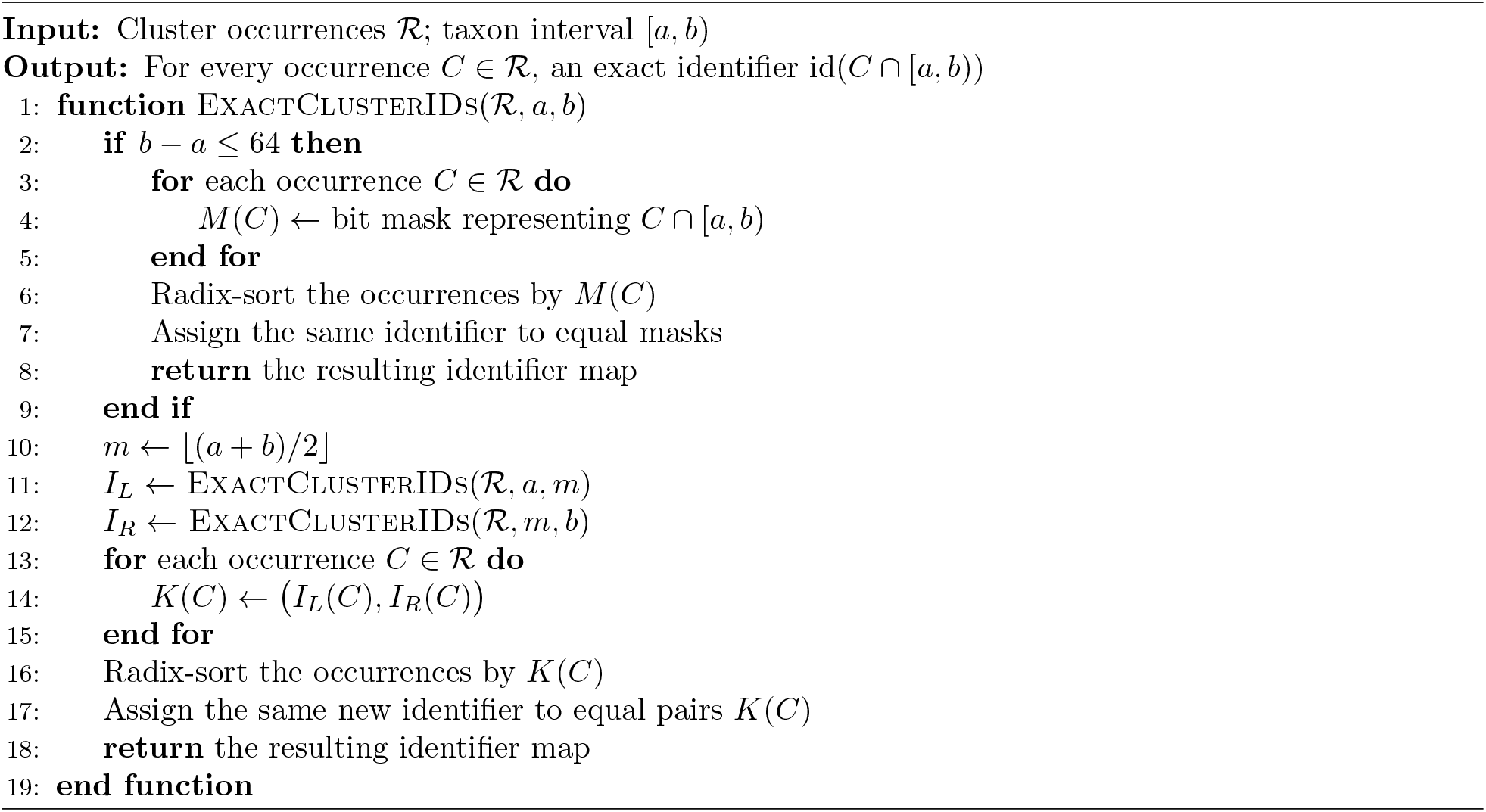

**Figure S1:**
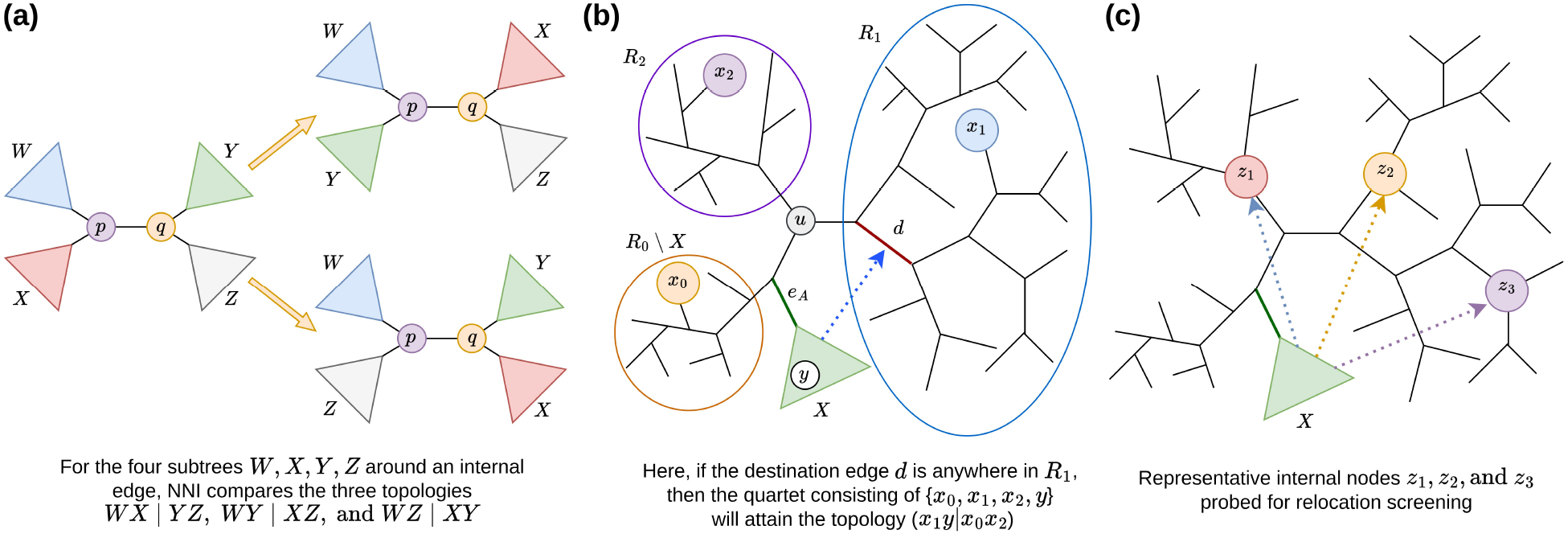
NNI and SPR refinement. (a) The three NNI resolutions around an internal edge. (b) For a fixed subtree *X*, the gain of reattaching it at an edge *d* is determined by local quartet contributions along the path from its current attachment. (c) A small set of representative internal nodes is used to screen promising relocation regions before the exact destination search.

#### S1.2.2 SPR refinement

NNI is effective at correcting local errors. However, moving a subtree over a long distance may often be helpful as well. To allow such longer-range corrections, QRAFT additionally uses SPR moves.

Let us consider a subtree *X* attached to the current tree through an edge *e_X_*. An SPR move cuts *X* from *e_X_* and reattaches it at another edge *d*. As in contextual regrafting, we would now like to find the destination edge that yields the largest increase in quartet support. However, there are Θ(*n*) possible subtrees and Θ(*n*) possible destination edges. Hence, an exhaustive search would be too costly.

We first observe that the score of moving a fixed subtree *X* has the same structure as the regrafting score of Sect. 2.5. Consider an internal node *u* outside *X*, and let *R*_0_, *R*_1_, *R*_2_ be the three subtrees incident to *u*, where *R*_0_ contains *X* (Fig. S1b). For *r* ∈ {0, 1, 2}, let *q_r_*(*u, X*) denote the number of gene-tree quartets containing one taxon from each of *X, R*_0_ \ *X, R*_1_, *R*_2_, in which the taxon from *X* is paired with the taxon from *R_r_*. These three quantities are obtained through the same four-color queries used for contextual regrafting. The only difference is in the assignment of the fourth color. In contextual regrafting, it represents the opposite side of the bipartition, whereas here the moving subtree *X* itself forms one color class and the remaining part of *R*_0_ forms another.

Now suppose that *X* is moved from its current attachment edge *e_X_* to another edge *d*. At any internal node outside the path between *e_X_* and *d*, both attachment edges lie in the same direction from that node, and its contribution therefore remains unchanged. Only the nodes along the path between the two attachment positions can affect the score difference. Consequently,

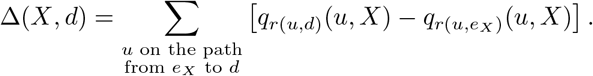

Thus, the gain of an SPR move can be obtained by accumulating local changes along the path from the current attachment to the proposed destination. Therefore, as in contextual regrafting, the quantities *q_r_*(*u, X*) are first computed for all internal nodes in *O*(*kn* log *n*) time. Then the gains for all destination edges can be obtained in *O*(*n*) time by traversing the tree and propagating the score thereby.

However, repeating this exact search for every possible subtree would lead to a quadratic search over the tree. We therefore use a heuristic screening step to identify a small number of promising subtrees, and perform the search only for those candidates.

We first restrict attention to candidate subtrees of sufficient size. For an edge *e*, let *w*(*e*) denote the number of taxa on its smaller side. We consider both sides of every edge with *w*(*e*) ≥ *m*_0_, where *m*_0_ is a fixed minimum size. Among the internal nodes of the tree, we select at most *z* representative nodes. For each such node *u*, we use its incident edges as sample candidate destinations. For every eligible subtree *X*, we compute the quartet-score gain of reattaching *X* to these edges and retain the best value (Fig. S1c). Importantly, the required gains for all eligible subtrees at a fixed *u* can be obtained together in *O*(*kn* log *n*) time using the four-color counting procedure with HDT traversal. Repeating the traversal over all *z* representative nodes therefore scores all eligible subtrees in *O*(*zkn* log *n*) time.

These estimated gains are used only for screening. We rank the candidate subtrees by their best estimated gain and retain at most the first *c* with positive gain. For each selected subtree, we then perform the exact search described above to determine its best destination over the full tree. The move is applied only if this exact calculation yields a positive quartet-score gain.

After a round in which at least one SPR move is accepted, we run NNI refinement again, since a longrange rearrangement may create new local improvements around the modified regions. We then repeat the SPR screening on the updated tree. The procedure performs at most *ρ* rounds and stops earlier if a round accepts no SPR move. Each round requires *O*(*zkn* log *n*) time for screening and at most *c* exact regrafting calls, each taking *O*(*kn* log *n*) time. Hence, the complete SPR refinement takes *O*(*ρ*(*z* + *c*)*kn* log *n*) time for fixed *z, c*, and *ρ*.

### S1.3 HDT-based Quartet Counting

QRAFT repeatedly requires us to solve the same underlying counting problem. Given a partition of the taxa into two or four color classes, we need to count how many quartets in the input gene trees show particular colored topologies. For example, when the two sides of a candidate bipartition are colored 0 and 1, the recursive divide needs the counts *W* (00 |11) and *W* (01 | 01). Contextual regrafting and NNI similarly require the three four-color counts *W* (01| 23), *W* (02 | 13), and *W* (03|12). We therefore need a way to compute these quantities efficiently.

A natural approach is dynamic programming. We can store suitable counting states on a tree and propagate them upward. Performing this computation directly on the gene trees, however, creates three difficulties. First, a gene tree may be highly unbalanced. Thus, changing the color of even a single leaf can require updating a linear number of ancestors in the worst case. Second, QRAFT needs families of closely related queries rather than single ones. For example, NNI refinement requires quartet support around every internal edge. Thus it is important to be able to move efficiently from one coloring of the taxa to the next. Finally, as the recursion progresses, a subproblem may involve only a small fraction of the original taxa. We thus need to restrict the computation to the active taxa without repeatedly processing the entire gene tree.

**Figure S2:**
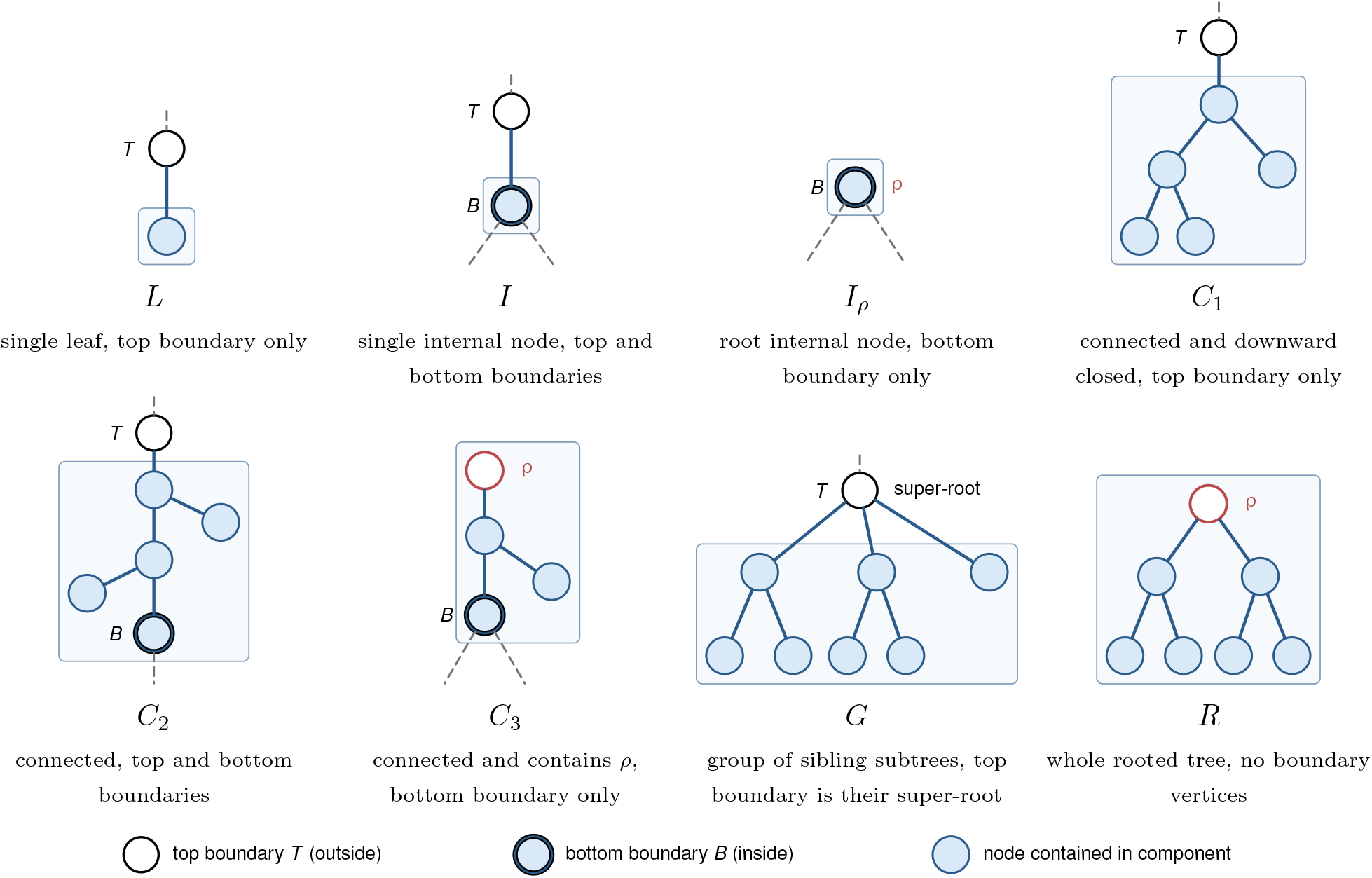
HDT component types. The four basic component types are L, I, C, and G. For counter bookkeeping, C is further divided into C_1_, C_2_, and C_3_, while R denotes the complete tree.

To address these challenges, we build on hierarchical decomposition trees (HDTs) [2–4]. An HDT reorganizes a gene tree into a locally balanced hierarchy of components. HDT also allows recoloring updates to be propagated through only a small part of the tree, while contraction and extraction let us process only the context relevant to the current subproblem. We now describe the HDT representation, the counters maintained at its components, the transition rules, and finally how a traversal of the query tree answers all required queries in quasi-linear time.

#### S1.3.1 Hierarchical decomposition trees

An HDT reorganizes a rooted tree into a locally balanced hierarchy of components [2, 3], where each component represents a set of nodes in the original tree. For a tree with *N* nodes, the HDT has *O*(*N*) size and can be constructed in *O*(*N*) time. A key property of HDT is local balance. If an HDT component represents a set of *s* nodes of the original tree, the subtree of the HDT rooted at that component has height *O*(log *s*). Thus, even when the original gene tree is highly unbalanced, its HDT provides a balanced structure on which updates can be propagated efficiently.

The HDT is built bottom-up with four standard types of components. An L component represents a single leaf of the gene tree, an I component a single internal node, a C component a connected set of nodes, and a G component a set of complete sibling subtrees. Initially, every gene-tree node forms an L or I component. These components are then repeatedly transformed and merged through the operations

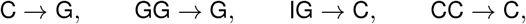

until one component represents the entire tree. Thus, the leaves of the resulting HDT correspond to the original gene-tree nodes (both leaves and internal nodes), while each internal node of the HDT represents the larger component formed by merging its child components. Fig. S2 illustrates the component types. Fig. S3 shows an example tree and its corresponding HDT. To define the counters below, we further divide the C components into three subtypes. A C_1_ component is downward closed and can connect to the rest of the tree only from above. A C_2_ component is open both upward and downward, while a C_3_ component contains the root and is open only downward. Finally, we use R to denote the closed component representing the entire tree.

**Figure S3:**
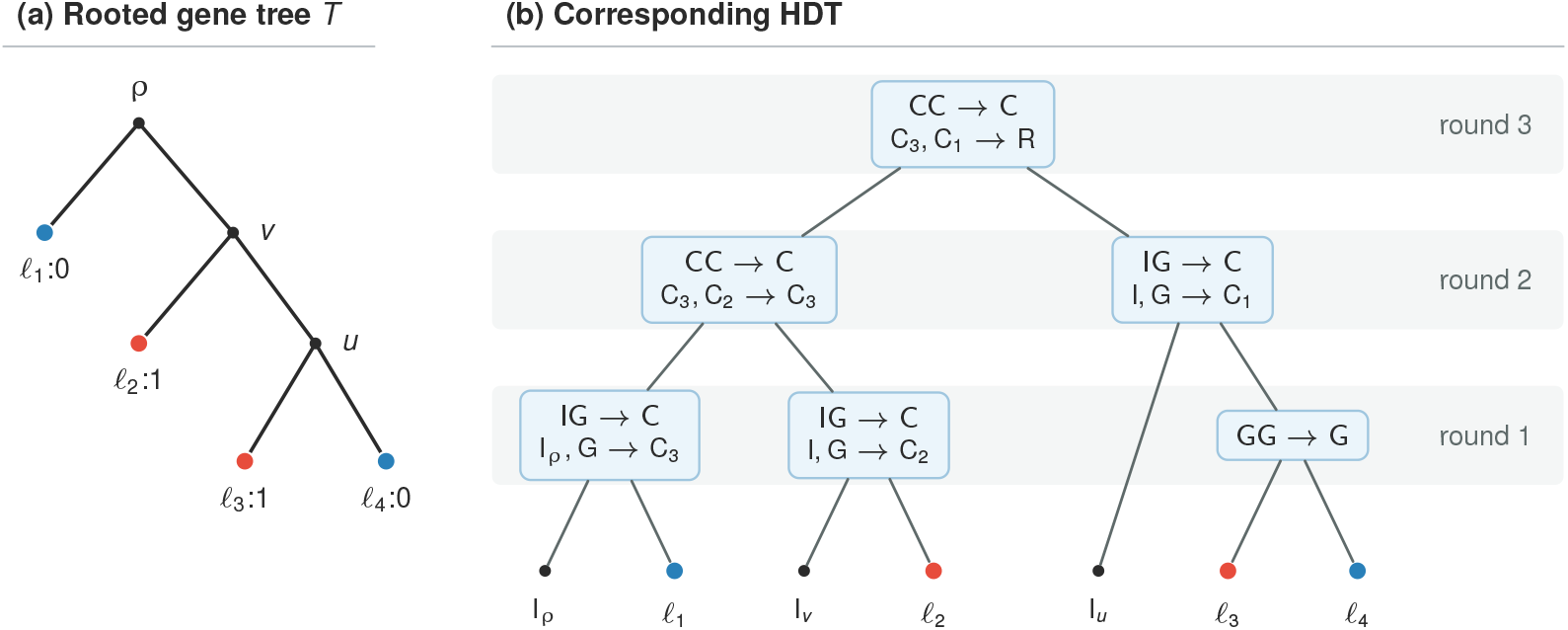
Example HDT construction. Leaves and internal vertices of *T* form the leaves of the HDT, while internal HDT nodes record successive component compositions. For each two-line annotation, the first line denotes the generic operation, while the second gives the specific component types involved.

We use the HDT as the data structure on which the colored quartet counts are maintained. Each component stores a constant number of counters summarizing the colored configurations inside it. The counters of a parent component are obtained from those of its children through transition equations. Thus, when the color of a leaf changes, we update its state and recompute only the HDT nodes on the path toward the root. The local balance property bounds this path by *O*(log *N*) nodes.

In fact, we often need to recolor a set of *r* taxa at once rather than a single taxon. Using the same local-balance property, the union of the corresponding *r* leaf-to-root paths can be updated in 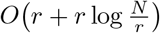 time [2], since nodes shared by multiple paths need to be recomputed only once. Thus, the time needed to recolor an entire subtree is nearly linear in the size of that subtree. We now consider the setting in QRAFT where we need to answer a family of related queries while traversing a query tree. For example, NNI requires the three quartet counts for every internal edge. To move efficiently from one query to the next, at each node we retain the coloring of its largest child and recolor only the remaining, smaller children. If a taxon belongs to such a smaller child, the size of the subtree containing it is at most half that of its parent. Hence, along any root-to-leaf path, the same taxon can be involved in such a recoloring only *O*(log *m*) times before the subtree size reaches one. Together with the path-union bound above, this gives *O*(*m* log *m*) total recoloring work for one gene tree over the complete traversal.

Finally, the recursive nature of QRAFT requires the data structure itself to shrink with the current sub-problem. Deep in the recursion, only a subset *S* of the original taxa may remain active,. Thus, retaining the complete gene-tree HDT would be unnecessarily expensive. We therefore use the Contractand Extract operations of Brodal et al. [3]. Given a selected set of active leaves, we first keep only the HDT paths needed to connect them to the root. Any region that contains no active leaf is then replaced by a single summarized component that stores the same counter information. We can further merge consecutive summarized regions and rebuild a locally balanced HDT containing only *O*(|*S*|) components. As a result, later recolorings and queries depend on the size of the current subproblem rather than on the size of the full gene tree, while producing the same quartet counts.

#### S1.3.2 Counters stored by an HDT component

We now specify the counter states stored at each HDT component. Our required outputs are quartet counts. This naturally leads to counters involving one to four leaves. Moreover, the counting tasks arising in QRAFT involve only a small set of quartet types. The two-color queries require 00 | 11 and 01 | 01, whereas the four-color queries require 01 | 23, 02 | 13, 03 | 12. Accordingly, we do not need to keep track of every possible configuration of colored leaves. It suffices to retain only those ones that can affect one of these five quartet counts after the component is combined with the rest of the tree.

**Table S1:**
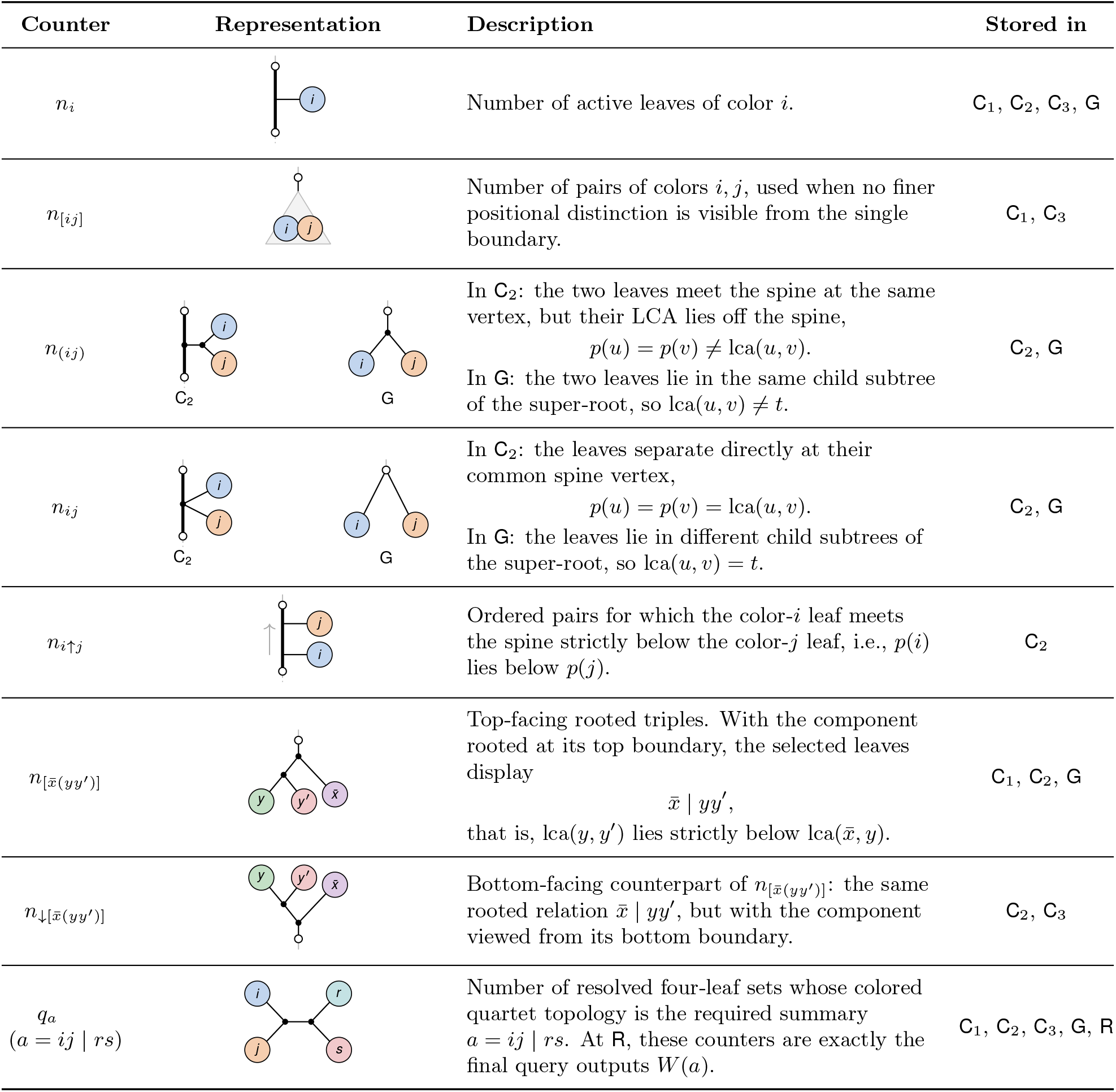
Counters stored at HDT components. Each row shows a counter, its represented configuration, and the corresponding component types. For C_2_, the vertical path denotes the spine between its two boundaries. Every non-root component also stores the implicit constant counter 1.

| Counter | Representation | Description | Stored in |
|---|---|---|---|
| $n_i$ |  | Number of active leaves of color $i$. | C<sub>1</sub>, C<sub>2</sub>, C<sub>3</sub>, G |
| $n_{[ij]}$ |  | Number of pairs of colors $i, j$, used when no finer positional distinction is visible from the single boundary. | C<sub>1</sub>, C<sub>3</sub> |
| $n_{(ij)}$ |  | In C<sub>2</sub>: the two leaves meet the spine at the same vertex, but their LCA lies off the spine,<br>$p(u) = p(v) \neq \mathrm{lca}(u, v)$.<br>In G: the two leaves lie in the same child subtree of the super-root, so $\mathrm{lca}(u, v) \neq t$. | C<sub>2</sub>, G |
| $n_{ij}$ |  | In C<sub>2</sub>: the leaves separate directly at their common spine vertex,<br>$p(u) = p(v) = \mathrm{lca}(u, v)$.<br>In G: the leaves lie in different child subtrees of the super-root, so $\mathrm{lca}(u, v) = t$. | C<sub>2</sub>, G |
| $n_{i\uparrow j}$ |  | Ordered pairs for which the color-$i$ leaf meets the spine strictly below the color-$j$ leaf, i.e., $p(i)$ lies below $p(j)$. | C<sub>2</sub> |
| $n_{[\bar{x}(yy')]}$ |  | Top-facing rooted triples. With the component rooted at its top boundary, the selected leaves display<br>$\bar{x} \mid yy'$,<br>that is, $\mathrm{lca}(y, y')$ lies strictly below $\mathrm{lca}(\bar{x}, y)$. | C<sub>1</sub>, C<sub>2</sub>, G |
| $n_{\downarrow[\bar{x}(yy')]}$ |  | Bottom-facing counterpart of $n_{[\bar{x}(yy')]}$: the same rooted relation $\bar{x} \mid yy'$, but with the component viewed from its bottom boundary. | C<sub>2</sub>, C<sub>3</sub> |
| $q_a$<br>$(a = ij \mid rs)$ |  | Number of resolved four-leaf sets whose colored quartet topology is the required summary $a = ij \mid rs$. At R, these counters are exactly the final query outputs $W(a)$. | C<sub>1</sub>, C<sub>2</sub>, C<sub>3</sub>, G, R |

Table S1 summarizes the resulting counters. The counter *n_i_* records the number of leaves of color *i*. For the single-boundary C components, C_1_ and C_3_, we use *n*_[*ij*]_ to simply count pairs of colors *i* and *j*, since their finer arrangement inside the component cannot be distinguished through the single exposed direction. The G and C_2_ components, however, require finer pair states. In G, these states denote whether the two leaves lie in the same or in different sibling subtrees. In C_2_, they instead describe how the leaves meet the path between its top and bottom boundaries. Let *π* denote this top-to-bottom path, which we call the spine. Let *p*(*u*) be the point where a leaf *u* meets *π*. The counters *n*_(*ij*)_, *n_ij_*, and *n_i↑j_* then distinguish whether two leaves enter through the same off-spine branch, separate directly at the same spine vertex, or meet the spine at different positions, respectively. Next, the triple counters 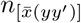 and 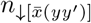 record the rooted relation 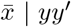 as viewed from the top and bottom boundaries, respectively. Finally, *q_a_* counts completed quartets of a required colored topology *a*.

The resulting counter vectors have the following sizes:

| Componenttype | *K* = 2 | *K* = 4 |
|---|---|---|
| C<sub>1</sub>, C<sub>3</sub> | 12 | 26 |
| C<sub>2</sub> | 23 | 56 |
| G | 15 | 32 |
| R | 2 | 3 |
| L | 3 | 5 |
| I, I<sub>*ρ*</sub> | 1 | 1 |

The largest state occurs for a four-color C_2_ component and contains only 56 counters. Thus, updating any HDT component requires only a small constant amount of arithmetic, independent of the number of taxa or the degree of the gene tree.

#### S1.3.3 Traversal and updates

The counter states defined above are computed bottom-up through two types of HDT transitions. The first are binary compositions, GG→G and CC→ C, which combine two child components into a larger one. The second are unary transformations, which change the component type without combining two independently represented leaf sets.

We first consider the binary case. Suppose that a new HDT component is formed by merging two smaller components. Any leaf set counted in the new component either lies entirely within one child or is split across the two. In the first case, the corresponding counter is carried forward directly. In the second, the new contribution is obtained by multiplying compatible counters from the two children. Thus, every parent counter can be written as

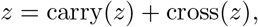

where carry(*z*) collects configurations already present in the children, while cross(*z*) counts those created by combining leaves from both children.

For example, when two G components are merged, the leaf counts simply add,

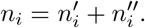

For a pair of colors *i, j*, a pair may already lie within either child, or its two leaves may come from different children. Hence,

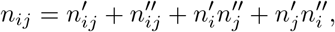

where coincident terms are counted only once when *i* = *j*.

The topology of the merge determines which pair counter receives the cross term. In a GG →G merge, leaves drawn from different children necessarily belong to different sibling subtrees and therefore contribute to *n_ij_*. In a CC →C merge, if the color-*j* leaf lies in the upper component and the color-*i* leaf in the lower component, the new pair instead contributes to *n_i↑j_* through the term 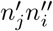.

The same principle extends to triples and quartets. A new triple spanning the two children must be formed by a 1 + 2 split of its leaves. A new quartet is formed either by a 1 + 3 split, combining a leaf with a compatible triple, or by a 2 + 2 split, combining two compatible pairs. Table S2 gives the resulting cross terms for the two binary HDT compositions, GG →G and CC →C. The full update of a counter is obtained by adding the corresponding counters already present in the children.

For compactness, let

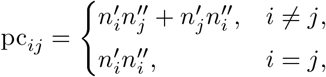

and, for GG → G, let

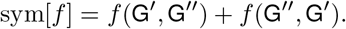

**Table S2:**
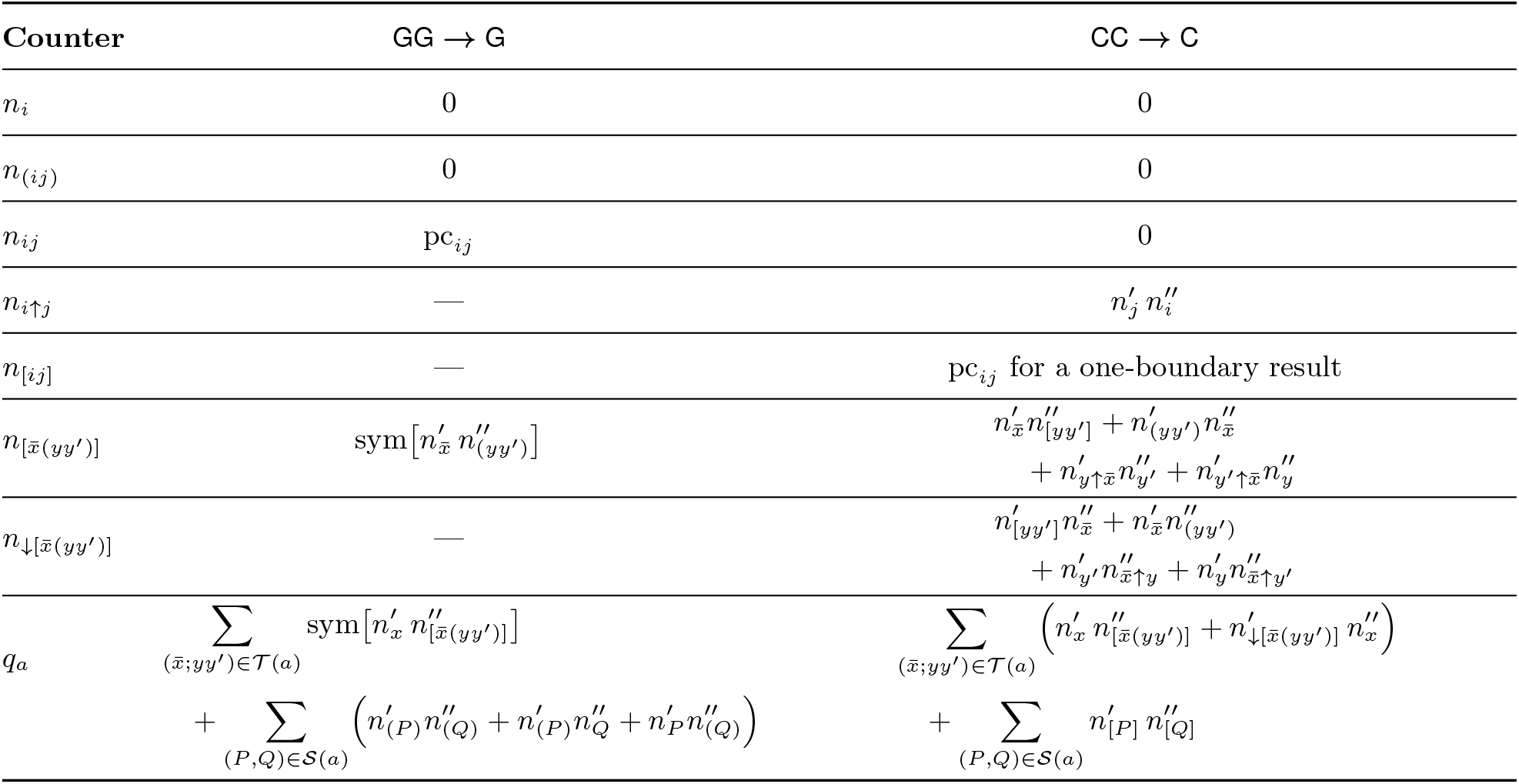
Counter transitions for the two binary HDT compositions. The table gives the cross term added to the counters inherited from the two children. For CC→ C, C′ and C″ denote the upper and lower components, respectively.

| **Counter** | GG → G | CC → C |
|---|---|---|
| *n*<sub>*i*</sub> | 0 | 0 |
| *n*<sub>(*ij*)</sub> | 0 | 0 |
| *n*<sub>*ij*</sub> | pc<sub>*ij*</sub> | 0 |
| *n*<sub>*i*↑*j*</sub> | — | $n'_j\, n''_i$ |
| *n*<sub>[*ij*]</sub> | — | pc<sub>*ij*</sub> for a one-boundary result |
| $n_{[\bar{x}(yy')]}$ | $\mathrm{sym}\big[n'_{\bar{x}}\, n''_{(yy')}\big]$ | $n'_{\bar{x}} n''_{[yy']} + n'_{(yy')} n''_{\bar{x}}$<br>$+ n'_{y\uparrow\bar{x}} n''_{y'} + n'_{y'\uparrow\bar{x}} n''_{y}$ |
| $n_{\downarrow[\bar{x}(yy')]}$ | — | $n'_{[yy']} n''_{\bar{x}} + n'_{\bar{x}} n''_{(yy')}$<br>$+ n'_{y'} n''_{\bar{x}\uparrow y} + n'_{y} n''_{\bar{x}\uparrow y'}$ |
| *q*<sub>*a*</sub> | $\sum_{(\bar{x};yy')\in\mathcal{T}(a)} \mathrm{sym}\big[n'_x\, n''_{[\bar{x}(yy')]}\big]$<br>$+ \sum_{(P,Q)\in\mathcal{S}(a)} \Big(n'_{(P)} n''_{(Q)} + n'_{(P)} n''_{Q} + n'_{P} n''_{(Q)}\Big)$ | $\sum_{(\bar{x};yy')\in\mathcal{T}(a)} \Big(n'_x\, n''_{[\bar{x}(yy')]} + n'_{\downarrow[\bar{x}(yy')]}\, n''_x\Big)$<br>$+ \sum_{(P,Q)\in\mathcal{S}(a)} n'_{[P]}\, n''_{[Q]}$ |

For a quartet channel *a*, let *T*(*a*) denote the rooted triples that can be completed to *a* by adding one appropriately colored leaf. For example, if *a* = 01| 23, then (1|23) ∈ *T* (*a*), since adding a leaf of color 0 gives 01 23, whereas (1 02) ∉ (*a*). Similarly, for a quartet channel *a* = *P*|*Q*, let *S*(*a*) denote the ordered assignments of the two color pairs to the two components. For example,

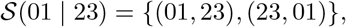

whereas (02, 13) ∉ *S*(01 | 23).

We also use

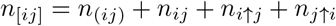

when only the undifferentiated number of pairs of colors *i, j* is required from a C_2_ component.

We next consider the unary transitions. These are simpler, since no selected leaf set is split across two components. A leaf of color *i* starts with *n_i_* = 1, while all higher-order counters are zero. The transformations C →G and IG → C mainly reinterpret the existing states under a new boundary structure. Their nontrivial updates are summarized in Table S3. Counters whose interpretation does not change are carried forward directly.

All of these transitions involve only the constant-size counter vectors described above. Therefore, once the states of the child components are known, the state of the resulting HDT component can be computed in constant time.

#### S1.3.4 Cost of quartet counting

We can now combine the locally balanced HDT with the traversal described above to obtain the quartet counting complexity used throughout QRAFT.

##### Theorem S1.1

(Cost of quartet counting). *For k single-copy gene trees and a query tree on m active taxa, one complete traversal computes all required quartet counts in O*(*km* log *m*) *time. Consequently, the quartet-counting work at a recursive subproblem on m taxa also takes O*(*km* log *m*) *time.*

**Table S3:** Initialization and boundary-changing transitions. Only counters whose interpretation changes are shown; the remaining compatible counters are carried forward unchanged.

| **Operation** | **Update** |
|---|---|
| Leaf initialization | For an active leaf of color $i$, we set $n_i = 1$, while all other nonconstant counters are zero. |
| C → G | A downward-closed C becomes one subtree below the super-root of G. Hence,<br>$n^{\mathsf{G}}_{(ij)} = n^{\mathsf{C}}_{[ij]}, \qquad n^{\mathsf{G}}_{ij} = 0.$ |
| IG → C<br>(two-boundary result) | The I node becomes the only new spine vertex, giving<br>$n^{\mathsf{C}}_{(ij)} = n^{\mathsf{G}}_{(ij)}, \qquad n^{\mathsf{C}}_{ij} = n^{\mathsf{G}}_{ij}, \qquad n^{\mathsf{C}}_{i\uparrow j} = 0,$<br>and<br>$n^{\mathsf{C}}_{\downarrow[\bar{x}(yy')]} = n^{\mathsf{G}}_{[\bar{x}(yy')]}.$ |
| IG → C<br>(one-boundary result) | When the finer pair placement is no longer visible,<br>$n^{\mathsf{C}}_{[ij]} = n^{\mathsf{G}}_{(ij)} + n^{\mathsf{G}}_{ij}.$<br>The corresponding root transformation produces C<sub>3</sub> and finally R in the same manner. |

*Proof.* Consider first a single gene tree. Since the two-color and four-color sets are fixed, every HDT component stores only a constant number of counters. Therefore, once the states of its children are known, recomputing the state of one component takes *O*(1) time.

Now suppose that a set of *r* taxa is recolored. Then only the union of their leaf-to-root paths in the HDT can change. For a locally balanced HDT with *N* nodes, Brodal et al. [2] show that this union contains

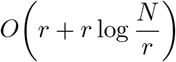

nodes. Hence, recoloring the entire set can be processed within the same bound by updating each affected HDT node once.

During the query-tree traversal, we retain the coloring of the largest child and recolor only the remaining, smaller children. Let *c* be such a child of a query-tree node *v*. The Extract and Contractoperations keep the current gene-tree HDT at size *O*(|*L*(*v*)|), so recoloring *L*(*c*) costs

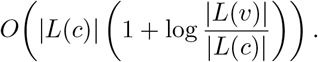

We charge this cost evenly to the taxa in *L*(*c*). Whenever a taxon receives such a charge, it lies in a child containing at most half as many taxa as its parent. Along any root-to-leaf path, the logarithmic ratios therefore telescope, and the total charge to one taxon is *O*(log *m*). Summing over all *m* taxa gives *O*(*m* log *m*) recoloring work for one gene tree. The temporary recolorings used by the four-color NNI queries follow the same smaller-child argument and do not change this bound.

The Extract operations have the same path-union cost, while successive Contractoperations along a largest-child path reduce the maintained structure geometrically [3]. Their total work is therefore absorbed by the same bound. Thus, one complete traversal costs *O*(*m* log *m*) per gene tree and

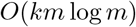

over all *k* gene trees.

At a recursive subproblem, QRAFT performs only a constant number of such traversals. Hence the total quartet-counting cost at that subproblem remains *O*(*km* log *m*).

### S1.4 Running Time and Memory

#### Theorem S1.2

(Running time and memory). *For k single-copy gene trees, a recursion of height h, a constant NNI move budget, and fixed SPR screening parameters, QRAFT runs in*

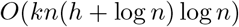

*time. Hence, the running time is O*(*kn* log^2^ *n*) *for a balanced recursion with h* = *O*(log *n*)*, and O*(*kn*^2^ log *n*) *in the worst case. With p workers, QRAFT uses O*(*kn* + *pn*) *memory.*

*Proof.* Consider a recursive subproblem on a taxon set *S* of size *m*. Since the gene trees are single-copy, each contains at most *m* leaves from *S*. After restriction and suppression of unnecessary degree-two vertices, each restricted gene tree has at most 2*m* nodes. Thus, the *k* restricted gene trees contain at most

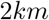

nodes in total.

QRAFT does not always materialize this exact restriction. Suppose instead that the current representation, of size *M*, is reused. Reuse occurs only when the exact restriction contains more than 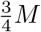 nodes. Since the restriction has at most 2*km* nodes,

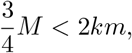

and therefore

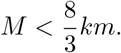

Hence, whether QRAFT keeps a new restriction or reuses the current representation, a subproblem on *m* taxa works with only *O*(*km*) gene-tree nodes.

By Theorem S1.1, quartet counting at this subproblem takes

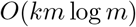

time. The remaining operations are no more expensive asymptotically. Restricting the MRE tree takes *O*(*m* log *n*) time, restricting the gene trees takes *O*(*km*), and the bipartition test requires *O*(*km* log *n*) time for a constant number of candidates and anchor pairs. Contextual regrafting uses the same quartet-counting machinery and remains within the same bound. Thus, the total work at a subproblem of size *m* is

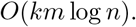

At any fixed recursion depth, the subproblems have disjoint taxon sets, so their sizes sum to at most *n*. Each depth therefore costs

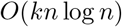

time. For recursion height *h*, the complete recursive divide and contextual regrafting stages require

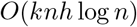

time.

We next account for refinement. In the default mode, QRAFT allows only a constant number of accepted NNI moves. Scoring all internal edges initially takes

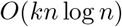

time. After an accepted move, only a constant number of nearby edges need to be rescored. Rescoring these edges still requires a pass over the gene trees and therefore takes *O*(*kn* log *n*) time in the worst case. Since the move budget is constant, all NNI scoring and rescoring together remains

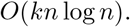

When compatible NNI moves are selected in a batch, the accompanying dynamic program takes *O*(*n* log *n*) time per pass and is dominated by the gene-tree processing.

For SPR refinement, let *z* be the number of representative internal nodes used for screening, *c* the maximum number of screened subtrees retained for exact evaluation, and *ρ* the maximum number of SPR rounds. In each round, screening all eligible subtrees at the *z* representative nodes takes

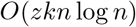

time. At most *c* selected subtrees then undergo an exact destination search, each taking

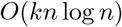

time. After a successful SPR round, QRAFT performs another bounded NNI refinement, again requiring *O*(*kn* log *n*) time. Thus, the complete refinement stage takes

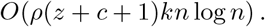

For fixed *z, c*, and *ρ*, together with a constant NNI move budget, this is

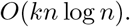

Finally, the MRE construction takes

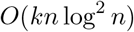

time. Combining the MRE construction, recursive inference, and refinement gives

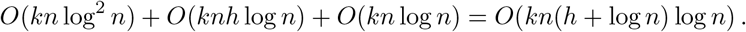

For a balanced recursion, *h* = *O*(log *n*), yielding

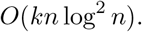

In the worst case, *h* = *O*(*n*), giving

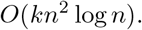

For memory, the input gene trees, the MRE tree, and their indexes require *O*(*kn*) space. Let *M*_0_ = *O*(*kn*) be the size of the representation at the root. Whenever a new restricted representation is retained, it contains at most ^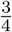^ as many nodes as its source. Thus, along any recursion path, the total size of the retained representations is bounded by

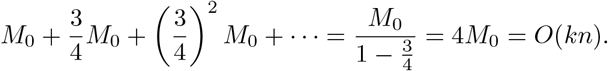

If a representation is reused, no additional full copy is created. Across simultaneously active subproblems, the active taxon sets are disjoint, and their counting and scoring structures therefore require *O*(*kn*) space in total. The NNI and SPR stages use only *O*(*n*) additional score, candidate, and dynamic-programming storage. Finally, each of the *p* workers requires *O*(*n*) temporary counting space. Hence, the total memory usage is

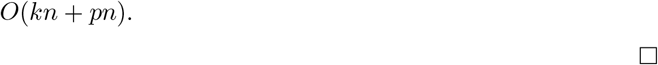

The running-time bound above applies to the bounded refinement used in the default configuration. In the extended mode, QRAFT removes the fixed NNI move budget, so its refinement time additionally depends on the number of accepted NNI moves and is not covered by Theorem S1.2. Empirically, however, none of the analyses reported in this study required more than 2,000 NNI passes.

## S2 Supplementary Results

### S2.1 Hardware

QRAFT’s running times come from three configurations. QRAFT used only the CPU, and ASTRAL-X and STELAR-X used the GPU:

- AMD EPYC 7742, 32 threads, 134 GB of RAM, NVIDIA RTX 4090 GPU
- AMD EPYC 9354, 64 threads, 439 GB of RAM, NVIDIA RTX 4090 GPU
- AMD EPYC 9354, 64 threads, 201 GB of RAM

We ran the other methods on an AMD EPYC 7742 with an RTX 4090 GPU, the runs also reported in the ASTRAL-X study [5]. The three-mode times on the biological datasets and ASTRAL-MP’s 10,000-taxon data, and the refinement-mode runs of Fig. S5, come from the EPYC 9354 configuration with 201 GB of RAM.

### S2.2 Scalability runs

For the comparisons with the other methods, QRAFT ran on one to five replicates per size. ASTRAL-X used tree-local search. ASTRAL-X and STELAR-X used GPUs, and ASTRAL-MP and ASTER had a 48-hour limit. The three-mode scaling curve (Fig. S5) is over the finished runs.

The extension of the STELAR-X dataset used 100,000–1,000,000 taxa with 1,000 gene trees and 100,000–1,000,000 gene trees with 1,000 taxa, in steps of 100,000, with five replicates per size and a 328 GB memory limit. We report means over the finished QRAFT runs, and ranges in the text.

QRAFT’s time on replicate 1 was at or near the top of its range. Three replicates were run at 600,000–800,000 taxa and at 700,000 to a million genes.

Accuracy on these datasets. With 1,000 true gene trees, QRAFT’s mean RF rate stayed between 0.39% and 0.56% at every size from 100,000 to one million taxa (0.46% at one million). On the replicates that ASTRAL-X and STELAR-X also completed, QRAFT’s RF rate was comparable or lower: 0.51% against 0.53% for both at 100,000 taxa (means over the same three replicates), and 0.57–0.58% against 0.62–0.64% at 200,000–400,000 taxa (replicate 1). On the STELAR-X datasets with up to 300,000 taxa or genes (Fig. S4), QRAFT’s mean RF rate was at most 0.02% above ASTRAL-X’s, and it was no higher from 100,000 taxa onward. With a million gene trees, QRAFT returned the true species tree on all three replicates.

**Figure S4:**
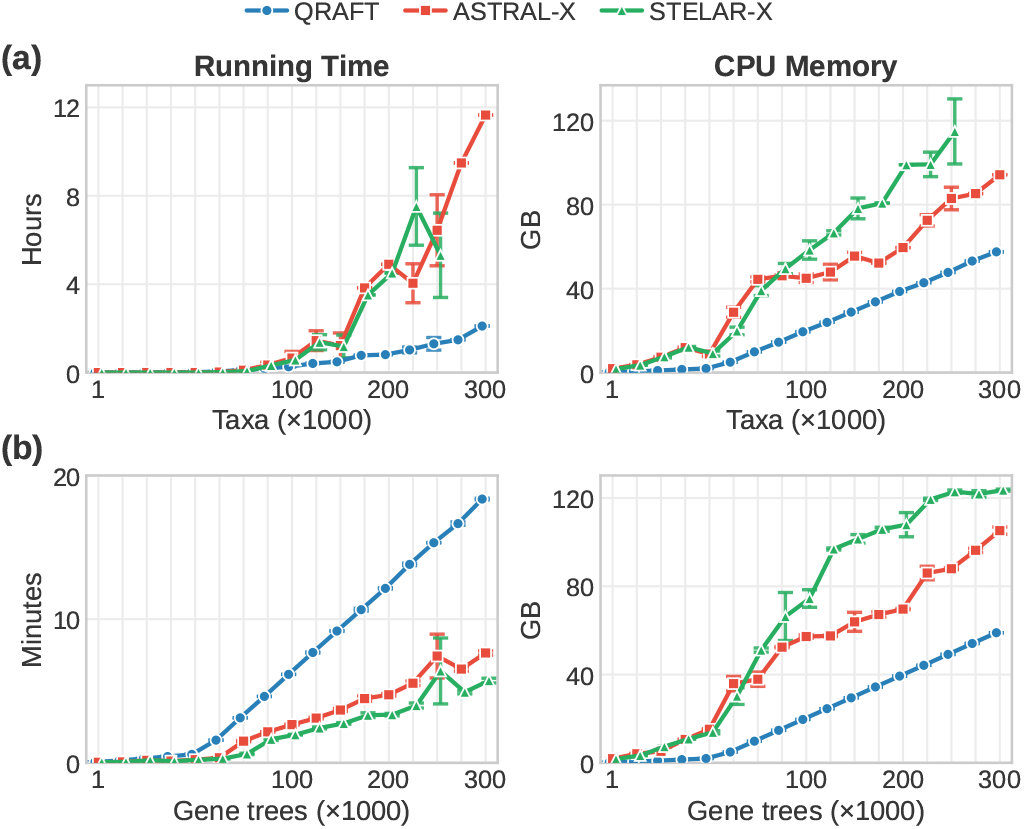
Average running time and peak CPU memory on the STELAR-X dataset: (a) 1,000 gene trees and 1,000–300,000 taxa, and (b) 1,000 taxa and 1,000–300,000 gene trees. Means over the one to five replicates per size that ASTRAL-X ran. STELAR-X was run up to 250,000 taxa, and both used their default search space S1. Time is in hours in (a) and minutes in (b).

### S2.3 Effect of the refinement mode

QRAFT has two refinement modes: default and extended. Default mode applies only a few NNI moves, while extended mode runs NNI optimization followed by up to two rounds of SPR refinement. We measured the running time and memory costs of both modes, as well as of no refinement at all, on the scalability datasets with medium ILS (ILS-L2, Fig. S5). Runtime was similar for default mode and no refinement, while extended mode increased runtime by a factor of 3–6.5×. At 100,000 taxa, no refinement, default mode and extended mode took 4.9, 5.4 and 35 minutes, respectively. At 100,000 gene trees, no refinement and default mode took 2.7 and 3.0 minutes, while extended mode took 11.4 minutes. It should be noted that in extended mode, NNI optimization phases took much less time compared to SPR, which took the majority of the time. SPR refinement constituted 1,793 seconds of the 2,108-second 100,000-taxon analysis, while NNI phases combined took 22 seconds. A similar trend (SPR: 514 seconds, NNI: 9 seconds, out of 686 seconds) was observed for 100,000 genes. All three modes grew close to linearly, as Fig. S5 shows. Peak memory also grew linearly. Default mode adds about a tenth to the time without refinement and suits fast routine analyses at scale. Extended mode takes about 4–5× as long on average with no extra memory and remains practical at 100,000 taxa (35 minutes) and beyond. Extended mode yields much higher quartet scores in some cases and is the mode we used on the biological datasets (Sect. 3.3).

### S2.4 Accuracy data and reference trees

The ASTRAL-II conditions have 1,000 estimated gene trees each. Their 200-taxon conditions have tree heights of 500K, 2M and 10M generations, which set high, medium and low ILS, and speciation rates 10^−6^ and 10^−7^. The 500- and 1,000-taxon conditions use 2M and 10^−6^. There are 47 and 50 replicates at high ILS with 10^−6^ and 10^−7^, respectively, 49 at 1,000 taxa and 50 else-where. The 10,000-taxon ASTRAL-MP dataset has nine replicates (R1–R10 without R9). The ablation study (Sect. S2.9) uses the ASTRAL-II and ASTRAL-MP datasets.

Fig. 5b uses the runs of the ablation study (Table S6): ASTRAL-X and STELAR-X in search space S3, ASTRAL-MP 5.15.5, ASTRAL-IV (ASTER v1.24.4.8) and the trees of the wQFM-TREE study [6]. Each 200-taxon bar averages the means of both speciation rates with equal weight. At 1,000 taxa, every method is shown on the 19 replicates that wQFM-TREE and ASTRAL-IV ran. ASTRAL-MP versions were 5.15.5 on the 37-taxon and ASTRAL-II datasets and 5.12.5 for its original study’s 10,000-taxon runs. The stored wQFM-TREE runs used for the 37-taxon true-gene-tree comparison came from its authors [6] and match ours on every replicate of the estimated conditions. STELAR-X needs rooted gene trees. As in the STELAR-X study, we rooted the estimated gene trees at the outgroup, and the 37-taxon true gene trees kept their simulated roots.

**Figure S5:**
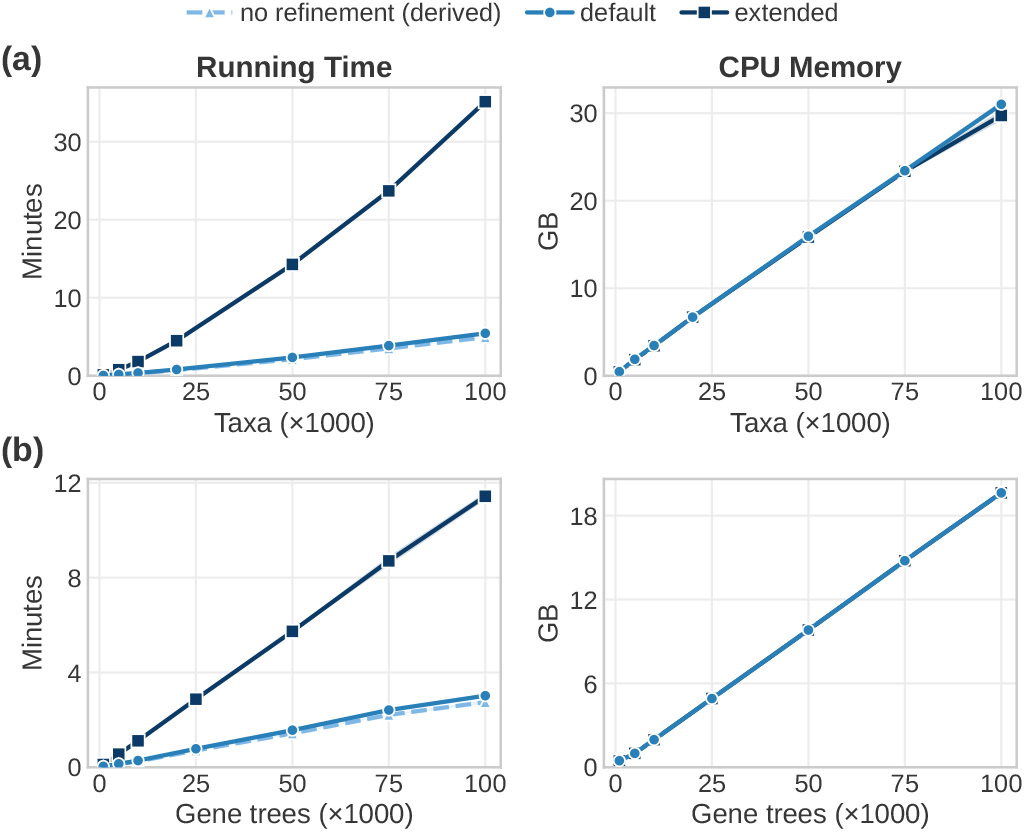
Mean running time and peak CPU memory of QRAFT with no refinement, in default mode, and in extended mode. (a) 1,000 gene trees and 1,000–100,000 taxa. (b) 1,000 taxa and 1,000–100,000 gene trees. Norefinement time was derived from default runs before NNI.

### S2.5 Biological comparisons

On the angiosperm dataset, uncapped NNI alone made 2,202 moves in 46 passes, and its tree scored 0.00007% below the published tree of Zuntini et al. [7]. In extended mode, NNI made 2,220 moves in 61 passes in all, and the SPR refinement made two moves in two rounds, reading 128 core vertices per round and calling the two-sided regraft four times. This raised the score to 0.00011% above theirs. All 23 bipartitions that the SPR refinement changed lie within Malpighiales: a Podostemaceae sample moved into its family’s clade, and the backbone around Euphorbiaceae, Achariaceae and Salicaceae and a few single-sample families was rearranged. The extended tree shares 8,496 of their tree’s 9,521 bipartitions, against 8,479 after NNI alone. At the level of orders, it differs from their tree in 22 of 142 bipartitions (default mode 40) and leaves 6 orders non-monophyletic (default mode 16, Zuntini et al. 2, ASTRAL-X 3).

For uDANCE, over all quartets resolved in the 380 gene trees (1.17×10^22^, scored with tqDist [4]), the normalized quartet score was 0.7273 for the default tree, 0.7344 for the published uDANCE tree of Balaban et al. [8] and 0.7345 for the extended NNI tree, which scored higher than the uDANCE tree on 203 of the 380 genes. Starting from that tree, the SPR refinement ran two rounds with 128 core-vertex reads each: 112 and 134 oracle calls applied 78 and 36 moves and raised the score to 0.7352 after the first round and 0.7353 after the second. Measured against GTDB, the uncapped-NNI tree had split Archaea (misplaced pieces of 258 and 7 tips), Patescibacteria (364 and 43), Negativicutes (268) and Melainabacteria (182 and 2). After the two rounds, each formed a single clade, and 101 GTDB phyla, 299 classes, 898 orders and 2,094 families were monophyletic. Spirochaetota stayed split (514 tips outside its largest clade). On the EPYC 9354 with 64 threads, the two rounds took 1.50 hours, and the refinement run peaked at 13.7 GB of memory.

**Table S4:** Running time and peak memory of QRAFT and other methods on the biological datasets. QRAFT was measured on an AMD EPYC 9354 with 64 threads. Its times are means of three runs, except the uDANCE 199k analysis, each run once. The other methods’ numbers are as reported by the ASTRAL-X study [5], measured on an AMD EPYC 7742 with 32 cores and an NVIDIA RTX 4090, so the times come from different hardware. ASTRAL-X, STELAR-X and ASTRAL-MP use the GPU, and QRAFT and ASTER do not. Guided uDANCE rows report single hierarchy steps. With the WoL1 step (86.8 s, 1.33 GB), the guided hierarchy took 3.80 h. The de novo 199k time comprises 4.05 h for the pipeline up to NNI and 1.50 h of SPR refinement. The angiosperm ASTRAL-MP time is from the original study’s run log, and ASTRAL-MP exceeded 129 GB when the ASTRAL-X authors reran it.

| Dataset (taxa; genes) | Method | Analysis | Time | CPU mem. (GB) | GPU mem. (GB) |
|---|---|---|---|---|---|
| uDANCE (199,330; 380) | QRAFT | de novo | 5.55 h | 14.05 | – |
|  | QRAFT | guided, 199k step | 3.71 h | 14.42 | – |
| uDANCE backbone (16,631; 380) | QRAFT | de novo | 282.7 s | 1.96 | – |
|  | QRAFT | guided, 16k step | 225.7 s | 2.03 | – |
| Angiosperms (9,524; 353) | QRAFT |  | 80.3 s | 1.33 | – |
|  | ASTRAL-X |  | 16 min | 34.6 | 3.0 |
|  | ASTRAL-MP |  | about 48 h | > 129 | – |

For the compute comparison in Sect. 3.3, we counted CPU hours from the marker alignments to the species tree. For uDANCE, we summed the CPU times of its 200K analysis in the data released by Balaban et al. [8]: 286,547 hours for the RAxML-NG gene trees and 18,465 hours for the IQ-TREE runs in its partitions, 770 hours for the APPLES-2 placement and 28,562 hours for ASTRAL, 334,344 hours in all. The analysis of the 16K backbone took another 44,896 hours, and the WoL tree that it extended is not counted. For QRAFT, we counted 1,180 CPU hours for the FastTreeMP gene trees and, for QRAFT itself, the wall-clock time multiplied by its 64 threads on the EPYC 9354, an upper bound on its CPU time: 355 hours *de novo* and 243 hours for the whole guided hierarchy. The two pipelines differ in their gene trees. uDANCE inferred maximum-likelihood gene trees with RAxML-NG within each partition, which took 86% of its CPU time, whereas we inferred one FastTree gene tree per marker. The CPU times were also measured on different hardware.

### S2.6 Major lineages in the uDANCE trees

#### Domain separation and CPR placement

QRAFT separated Archaea and Bacteria exactly, consistent with Balaban et al. [8]. It placed the Candidate Phyla Radiation (CPR, called Patescibacteria in GTDB) as the sister of all other Bacteria, consistent with Zhu et al. [9] and Balaban et al. [8]. Hug et al. [10] described CPR as a major bacterial radiation. Zhu et al. noted that long-branch attraction could cause the basal placement. Coleman et al. [11] and Martinez-Gutierrez and Aylward [12] instead placed CPR within Terrabacteria as the sister of Chloroflexota. Terrabacteria is the bacterial superphylum that includes Actinobacteriota, Firmicutes, Cyanobacteria and Chloroflexota [13].

#### Terrabacteria clade, the DST phyla and Fusobacteriota

QRAFT recovered Terrabacteria as one clade (76,219 genomes), apart from 131 genomes de novo and 138 guided. QRAFT placed the DST phyla (Deinococcota, Synergistota and Thermotogota) and Fusobacteriota within Terrabacteria. For Deinococcota, this is consistent with Battistuzzi et al. [13] and Balaban et al. [8]. Coleman et al. [11] also placed the DST phyla on the Terrabacteria side, but could not resolve where Fusobacteriota attaches. Martinez-Gutierrez and Aylward [12] instead placed Thermotogota and Synergistota at the base of Bacteria. Balaban et al. [8] placed Fusobacteriota, Synergistota and Thermotogota next to Gracilicutes, the other major bacterial clade, which includes Proteobacteria, the FCB group and the PVC group [11]. Within Terrabacteria, QRAFT placed Fusobacteriota as the sister of Firmicutes, a relationship that earlier rRNA and ribosomal-protein trees suggested [14]. Balaban et al. [8] instead placed Fusobacteriota as the sister of Thermotogota and its relatives.

#### QRAFT recovered Proteobacteria and its two large classes

Several large groups that the published uDANCE tree breaks were monophyletic in QRAFT. In the guided tree, Proteobacteria (61,040 genomes) was monophyletic, whereas Balaban et al. [8] noted that a handful of genomes with placeholder names break it in their tree. QRAFT recovered Gammaproteobacteria (48,898 genomes), which in GTDB includes the former Betaproteobacteria as the order Burkholderiales [15]. The uDANCE tree missed it by one genome. The guided tree also recovered Alphaproteobacteria (12,075 genomes), which the uDANCE tree missed by 14 genomes.

#### QRAFT recovered Planctomycetota, Paceibacteria and SAR86

QRAFT recovered Planctomycetota, which the uDANCE tree missed by six of its 822 genomes. QRAFT also recovered the CPR class Paceibacteria, which the uDANCE tree missed by two of its 1,150 genomes. QRAFT kept all 440 genomes of the order SAR86 together with high support (100%), whereas the uDANCE tree split the order into two clades of 351 and 89 genomes. QRAFT, however, grouped Chlamydiota with the Spirochaetota class Leptospirae, outside the PVC group (Planctomycetota, Verrucomicrobiota and Chlamydiota). Balaban et al. [8] instead kept Chlamydiota in PVC, as Zhu et al. [9] did, although Zhu et al. found the placement of Chlamydiae sensitive to taxon sampling.

#### QRAFT recovered Thermoplasmatota, Bacteroidales and Bipolaricaulota

Guided QRAFT recovered Thermoplasmatota (882 genomes) and the order Bacteroidales (5,814 genomes), which the uDANCE tree missed by two genomes and one genome, respectively. In QRAFT, the FCB group of the NCBI taxonomy (Fibrobacterota, Chlorobiota, Bacteroidota and related phyla), to which Bacteroidales belongs, was about as consistent with that taxonomy as in the uDANCE tree. De novo, all 16 Bipolaricaulota genomes formed one clade with high support (100%), whereas Balaban et al. [8] found this phylum split into distant subclades, and their tree misplaced 9 of its genomes. QRAFT de novo also grouped the GTDB phyla Myxococcota and Myxococcota A together, with 2 of their combined 350 genomes misplaced, against 46 in the uDANCE tree. Neither phylum was monophyletic on its own in QRAFT, consistent with Balaban et al. [8], who noted that Myx-ococcota may be truly paraphyletic. De novo, QRAFT also recovered Chloroflexota, which the uDANCE tree missed by one genome. Guided QRAFT instead placed the Chloroflexota class Limnocylin-dria with Dormibacterota, the sister phylum of Chloroflexota [11].

#### Placement of Thermotogota and its relatives

QRAFT placed Dictyoglomota, Thermotogota, Caldisericota and Coprothermobacterota away from the base of non-CPR Bacteria, consistent with the final uDANCE tree of Balaban et al. [8]. QRAFT grouped the four phyla with Synergistota. Balaban et al. [8] instead grouped them with Fusobacteriota, with low support (3%).

**Figure S6:**
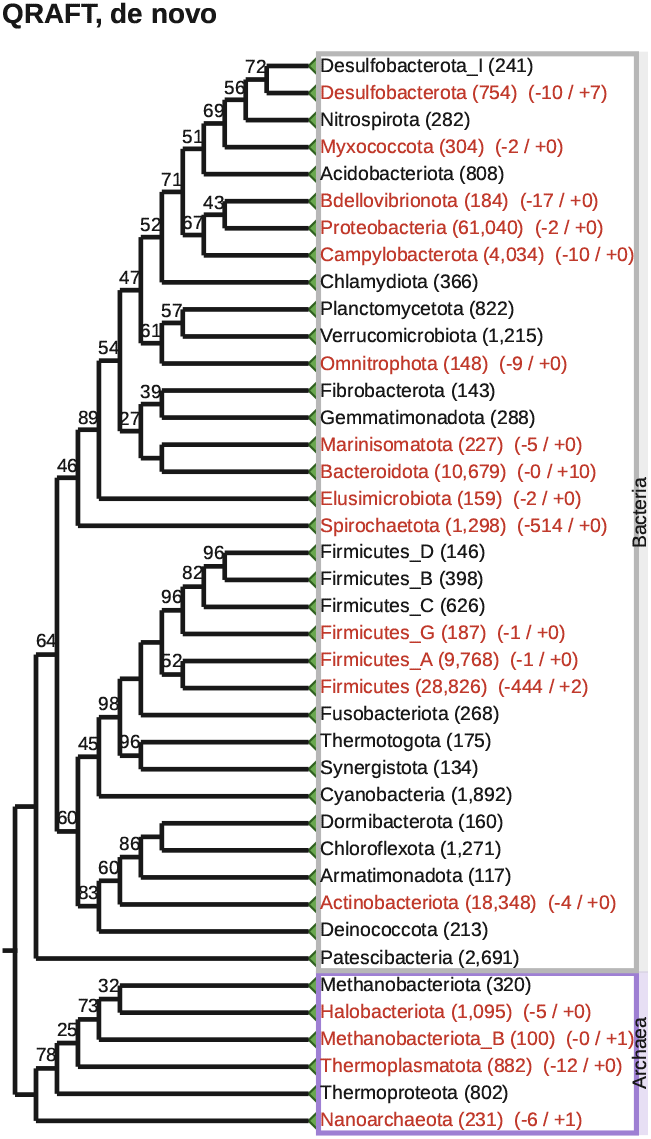
QRAFT’s de novo uDANCE tree, rooted on Archaea and drawn as in Fig. 6. The 40 GTDB phyla with at least 100 genomes are collapsed, and 22 of them are monophyletic.

#### Placement of DPANN within Archaea

QRAFT placed DPANN (Diapherotrites, Parvarchaeota, Aenigmarchaeota, Nanoarchaeota, Nanohaloarchaeota and related lineages) as the earliest-diverging archaeal lineage, consistent with Balaban et al. [8]. Williams et al. [16] and Dombrowski et al. [17] also suggested this placement, although Martinez-Gutierrez and Aylward [12] questioned it as a possible artifact. QRAFT also grouped Asgard with the TACK superphylum (Thaumarchaeota, Aigarchaeota, Crenarchaeota and Korarchaeota), which GTDB treats as the phylum Thermoproteota [18], consistent with Martinez-Gutierrez and Aylward [12].

**Table S5:**
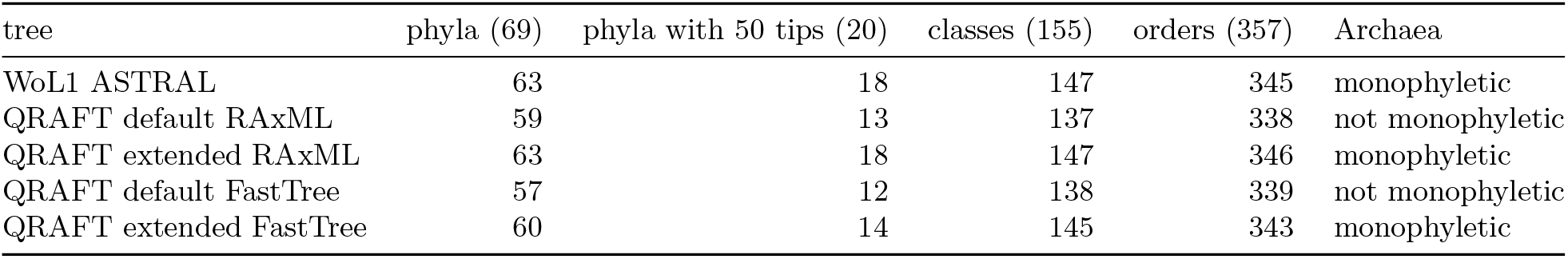
Monophyletic GTDB groups on the WoL1 genomes, with RAxML and FastTree gene trees of the same 380 markers.

#### Groups that QRAFT splits

Some GTDB groups recovered by uDANCE were not monophyletic in QRAFT’s trees. In some of these disagreements, NCBI agrees with QRAFT. Notably, QRAFT separates 50 genomes (59 *de novo*) of the order Mycobacteriales, mostly of the family Frankiaceae, which NCBI places in a separate order, Frankiales. Similarly, QRAFT separates 224 genomes of the order Peptostreptococcales, most of which NCBI places in a separate order, Eubacteriales. There are also cases such as the class Desulfurellia (10 genomes), which QRAFT separates from Campylobacterota although both GTDB and NCBI place it in that phylum. Still, moving any of these clades back into its group lowers the exact quartet score of both QRAFT trees. The differences therefore come from the signal in the gene trees, not from a failure of QRAFT’s search, and may reflect genuine alternative relationships, non-tree-like evolution or gene-tree estimation error, as discussed in the main text.

### S2.7 WoL1 control: FastTree versus RAxML gene trees

We compared QRAFT results from two gene-tree inputs for 380 markers on the 10,575 WoL1 genomes. FastTree gene trees inferred from the uDANCE alignments were pruned to these genomes and matched to WoL1’s own RAxML marker trees by normalized Robinson–Foulds (RF) distance on common genomes. All 380 markers matched one to one. We ran QRAFT in default and extended modes on each input.

The per-gene normalized RF distance between FastTree and RAxML trees had a median of 0.264, an interquartile range of 0.239–0.291, and a range of 0.174–0.390. We assessed monophyly using GTDB r207 labels for 9,698 tips and groups with at least two tips. Table S5 reports the numbers of monophyletic groups. With RAxML gene trees, QRAFT extended matched the WoL1 ASTRAL tree in the numbers of mono-phyletic phyla and classes. With FastTree gene trees of the same markers, it recovered 3 fewer phyla, 2 fewer classes and 3 fewer orders than its RAxML run. Extended mode made Archaea monophyletic on both inputs. However, the FastTree trees came from the uDANCE (WoL2) alignments and the RAxML trees from the WoL1 alignments. Thus, the per-gene differences mixed tree method and alignment effects.

### S2.8 Angiosperm datasets

#### S2.8.1 Analysis of the angiosperm dataset

We reanalyzed the angiosperm dataset of Zuntini et al. [7], containing 9,524 taxa and 353 gene trees with missing taxa. QRAFT reconstructed the angiosperm phylogeny in only 80.3 seconds, using only 1.33 GB of CPU RAM. For comparison, ASTRAL-X needed 16 minutes and 34.6 GB of CPU RAM on this dataset, and the ASTRAL-MP run of the original study took about 48 hours [5]. Importantly, the species tree inferred by QRAFT (Fig. S7) recovered the major relationships of the published angiosperm tree of Zuntini et al. [7] and was broadly consistent with earlier phylogenomic work [19]. Its normalized quartet score was 0.852756, 0.00011% above that of Zuntini et al. (0.852755).

To further assess the quality of the inferred tree, we examined whether each recognized angiosperm order forms a monophyletic clade in the QRAFT species tree. Among the 74 recognized orders represented in the dataset, 68 (91.9%) were recovered as monophyletic without any post-processing. Each of the remaining six orders differed from monophyly by a very small number of taxa separated from an otherwise intact clade, affecting a total of only 10 taxa across the dataset. Arecales (separated by its four Dasypogonaceae taxa) and Escalloniales (separated by two taxa) are not monophyletic, consistent with Zuntini et al. [7]. Alismatales, Poales, Lamiales and Malpighiales are monophyletic in Zuntini et al. [7]. In the QRAFT tree, three taxa with low gene coverage break them: *Eminium spiculatum* (237 genes), which QRAFT moved from Alismatales into Poales, *Pycnanthemum tenuifolium* (145 genes, Lamiales) and *Pimelodendron amboinicum* (167 genes, Malpighiales). Zuntini et al.’s placements of these three taxa have support near zero on these gene trees. The quartet objective itself favors QRAFT’s placement of *Pycnanthemum* and *Pimelodendron*: moving either back to where Zuntini et al. placed it lowers the quartet score.

**Figure S7:**
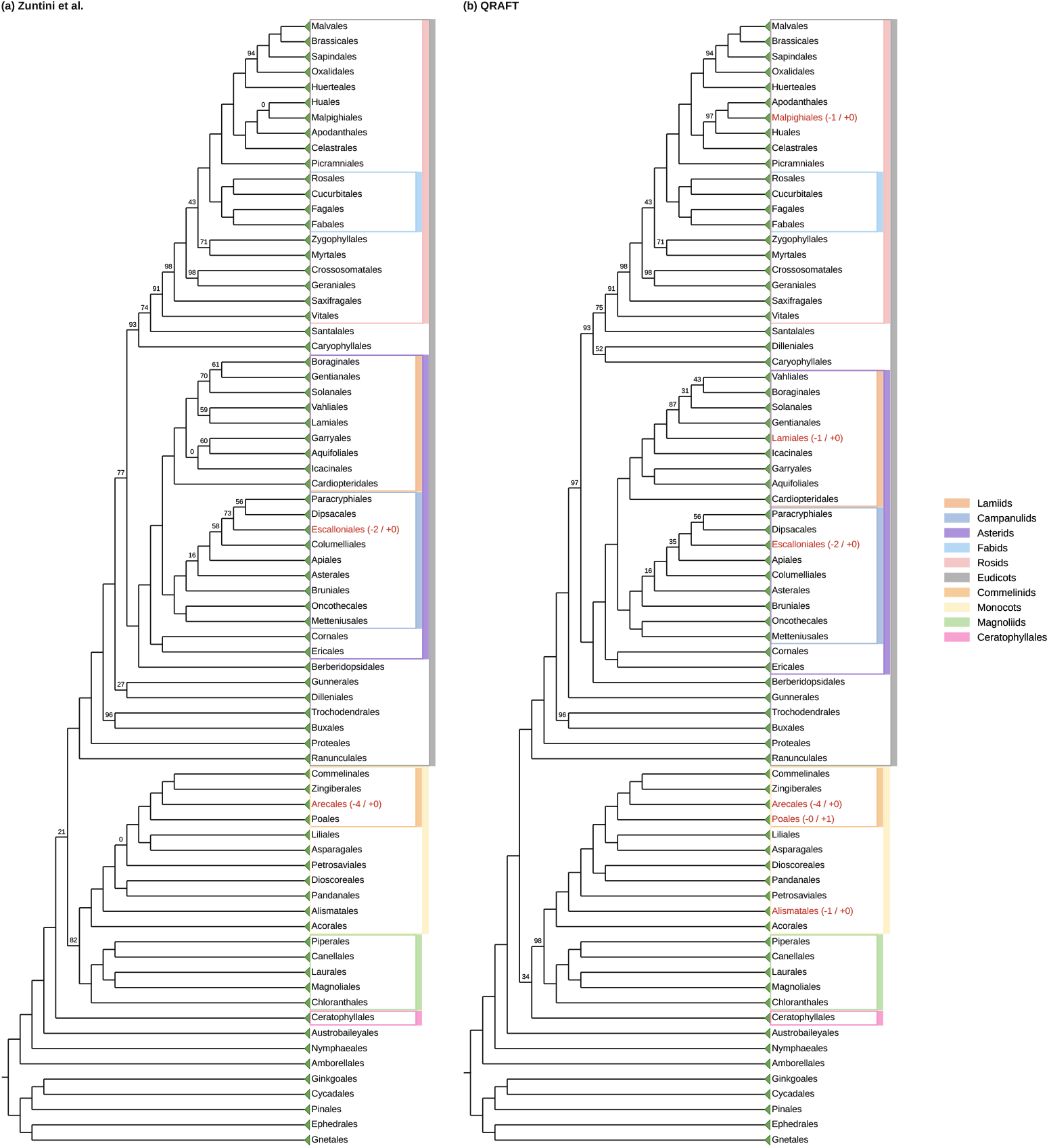
Angiosperm trees: (a) the published tree of Zuntini et al. [7] and (b) QRAFT, rooted on the gymnosperm orders, with orders collapsed and major clades marked.

We next present several observations on the angiosperm phylogeny inferred by QRAFT in comparison with previous studies.

#### QRAFT recovered relationships among Asterids

Consistent with Zuntini et al. [7], QRAFT recovered the Asterid relationships. Apart from *Pycnanthemum tenuifolium*, Ericales and Cornales formed a clade sister to the remaining Asterids, as in the 1KP study [19]. The other orders were divided into two major clades, Lamiids and Campanulids. Within these clades, Metteniusales and Oncothecales were sister to all other Campanulids, and Cardiopteridales was sister to all other Lamiids, both consistent with Zuntini et al.

#### Placement of Amborellales and Ceratophyllales

QRAFT placed Amborellales as sister to all other angiosperms, consistent with Zuntini et al. [7]. It placed Ceratophyllales sister to Monocots, Magnoliids and Chloranthales with low support (0.34). The placement of Ceratophyllales has also conflicted among earlier studies [19, 20].

#### QRAFT recovered the nitrogen-fixing clade

Within the Rosids, QRAFT grouped Cucurbitales, Fabales, Fagales and Rosales into one clade, as in Zuntini et al. [7]. This clade has long been supported by evidence of a single origin of the predisposition for symbiotic nitrogen fixation [7, 21].

#### Placement of Oxalidales

Oxalidales grouped with Malvales, Brassicales and Sapindales. QRAFT recovered this relationship with support 0.94, as did Zuntini et al. [7], rather than the Celastrales–Oxalidales– Malpighiales clade of APG IV [22].

#### Superrosids and major angiosperm clades

QRAFT recovered Superrosids (support 0.91), consistent with Zuntini et al. [7]. ASTRAL-X separated the 22 Vitales taxa from this clade. Monocots, Magnoliids, Magnoliids with Chloranthales, Eudicots and core Eudicots were all recovered.

#### Weakly supported placements consistent with whole-genome duplication and early hybridization

QRAFT placed Dilleniales sister to Caryophyllales with low support (0.52). This placement differs from Zuntini et al. [7], who placed Dilleniales with Gunnerales. The low support is consistent with ancient whole-genome duplications in these lineages [19, 23]. The branch that joins Ceratophyllales to Monocots, Magnoliids and Chloranthales, and so separates them from the Eudicots, also received low support (0.34). This is consistent with early hybridization among the Mesangiosperm lineages [7, 24].

### S2.9 Ablation study

QRAFT breaks the problem of species tree inference into a handful of steps: starting tree construction, divide score, bipartition test and refinement. To validate these steps and assess the importance of each, we evaluated QRAFT with variants of each step or with each step turned off (Table S6). The study used the eight model conditions of ASTRAL-II and the 10,000-taxon dataset of ASTRAL-MP, described with the accuracy datasets in Sect. 3.

We study the following variants of QRAFT’s pipeline before refinement, each followed by no, default or extended refinement: (i) the MRE tree only, (ii) the refined MRE tree, (iii) the balanced divide and (iv) no bipartition test. For (i), we skipped the divide and the regrafting. For (ii), QRAFT started from the MRE tree after NNI and SPR refinement in extended mode, which raises its quartet score. Since the MRE tree may be unresolved, refinement on it moves only at vertices of degree three. Scoring moves at vertices of higher degree would need counting with more colors, as for trees of arbitrary degree in Brodal et al. [3], which we do not implement. For (iii), instead of selecting and testing the MRE edge with the highest score *s*(*A* | *B*), QRAFT selected and tested the edge that divides the subproblem’s taxa most evenly. For (iv), we skipped the bipartition test. We used the same versions of ASTRAL-MP, ASTRAL-X, STELAR-X and wQFM-TREE as in the accuracy analyses (Sect. 3.2). All quartet-score changes Δ*q* are relative to QRAFT’s default tree. ASTRAL-IV was not run on ASTRAL-MP’s 10,000-taxon dataset, as it did not finish 10,000 taxa within 48 hours even with true gene trees (Fig. 5c).

#### QRAFT’s default pipeline

QRAFT’s default pipeline was chosen because it is well balanced (Table S6). On every ASTRAL-II column it was the most accurate variant or close to it, with an RF rate at most 0.16% higher. Refinement mattered mainly under high ILS, where default NNI lowered the RF rate from 6.92% to 5.77%. Elsewhere it lowered the RF rate by at most 0.21%. Extended mode raised the quartet score in every column, but it lowered the RF rate only under high ILS and raised it in the other columns. Its changes were mostly NNI moves: on these conditions, SPR moves with a positive gain were rare (107 SPR moves against 5,495 NNI moves over all 375 runs). A higher quartet score does not mean a more accurate tree, as the quartet score may overfit to gene-tree estimation error. Indeed, QRAFT’s default tree already scored above the true tree on most replicates. QRAFT therefore keeps capped, gated NNI as its default, with extended mode as an option where maximizing quartet support is the objective. Default refinement adds only about a tenth to the running time without refinement, whereas extended mode takes about 4–5× as long (Fig. S5).

**Table S6:**
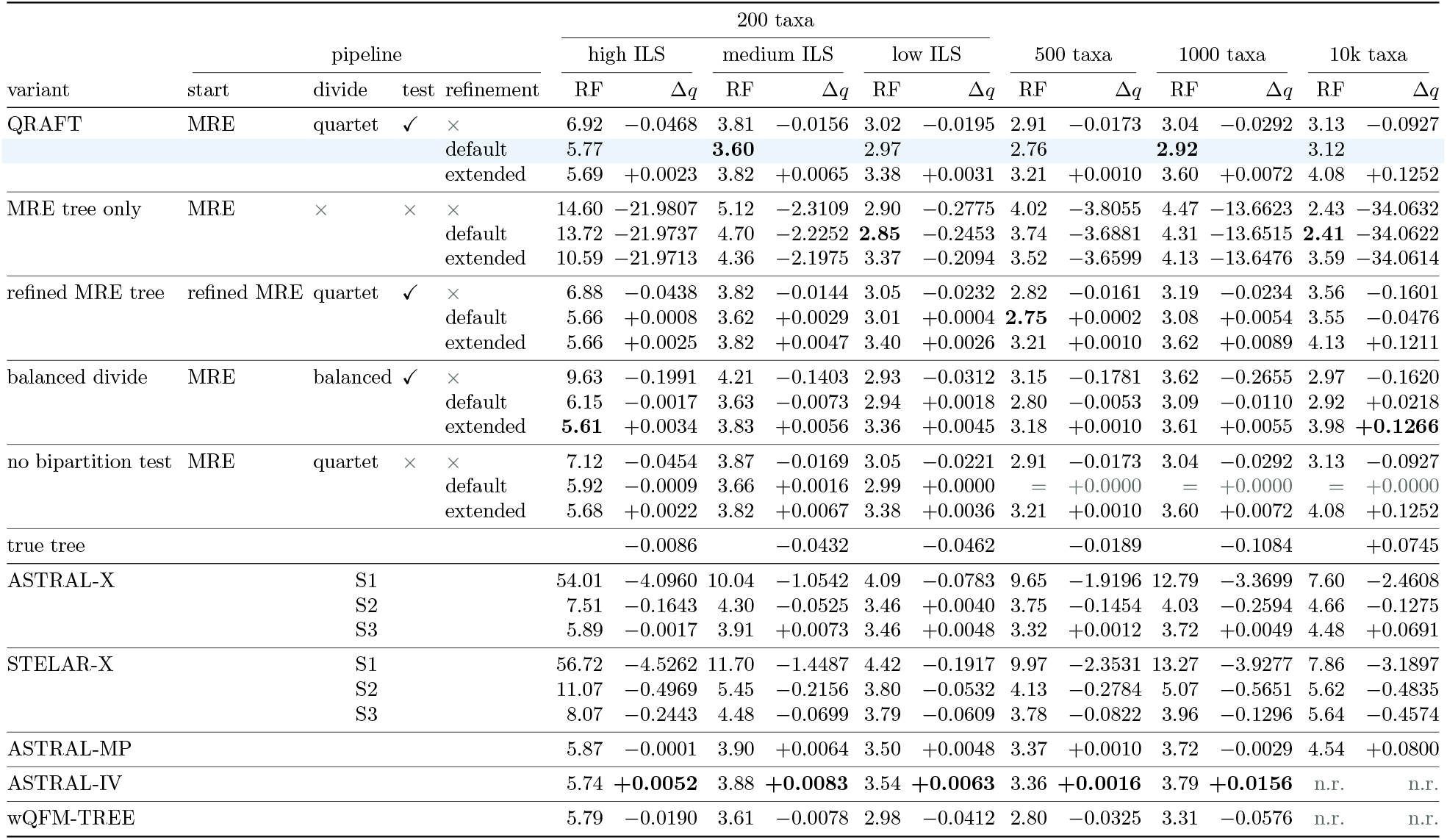
Mean RF rate (%) and mean quartet-score change Δ*q* from QRAFT’s default tree (% of genetree quartets) on ASTRAL-II’s eight model conditions and ASTRAL-MP’s 10,000-taxon dataset with 1,000 estimated gene trees. Symbols: × step not used, = QRAFT’s default rate on every replicate, gray no score change on any replicate, n.r. not run.

|  |  |  |  |  | 200 taxa |  |  |  |  |  |  |  |  |  |  |  |
|---|---|---|---|---|---|---|---|---|---|---|---|---|---|---|---|---|
|  | pipeline |  |  |  | high ILS |  | medium ILS |  | low ILS |  | 500 taxa |  | 1000 taxa |  | 10k taxa |  |
| variant | start | divide | test | refinement | RF | Δ*q* | RF | Δ*q* | RF | Δ*q* | RF | Δ*q* | RF | Δ*q* | RF | Δ*q* |
| QRAFT | MRE | quartet | ✓ | × | 6.92 | −0.0468 | 3.81 | −0.0156 | 3.02 | −0.0195 | 2.91 | −0.0173 | 3.04 | −0.0292 | 3.13 | −0.0927 |
|  |  |  |  | default | 5.77 |  | **3.60** |  | 2.97 |  | 2.76 |  | **2.92** |  | 3.12 |  |
|  |  |  |  | extended | 5.69 | +0.0023 | 3.82 | +0.0065 | 3.38 | +0.0031 | 3.21 | +0.0010 | 3.60 | +0.0072 | 4.08 | +0.1252 |
| MRE tree only | MRE | × | × | × | 14.60 | −21.9807 | 5.12 | −2.3109 | 2.90 | −0.2775 | 4.02 | −3.8055 | 4.47 | −13.6623 | 2.43 | −34.0632 |
|  |  |  |  | default | 13.72 | −21.9737 | 4.70 | −2.2252 | **2.85** | −0.2453 | 3.74 | −3.6881 | 4.31 | −13.6515 | **2.41** | −34.0622 |
|  |  |  |  | extended | 10.59 | −21.9713 | 4.36 | −2.1975 | 3.37 | −0.2094 | 3.52 | −3.6599 | 4.13 | −13.6476 | 3.59 | −34.0614 |
| refined MRE tree | refined MRE | quartet | ✓ | × | 6.88 | −0.0438 | 3.82 | −0.0144 | 3.05 | −0.0232 | 2.82 | −0.0161 | 3.19 | −0.0234 | 3.56 | −0.1601 |
|  |  |  |  | default | 5.66 | +0.0008 | 3.62 | +0.0029 | 3.01 | +0.0004 | **2.75** | +0.0002 | 3.08 | +0.0054 | 3.55 | −0.0476 |
|  |  |  |  | extended | 5.66 | +0.0025 | 3.82 | +0.0047 | 3.40 | +0.0026 | 3.21 | +0.0010 | 3.62 | +0.0089 | 4.13 | +0.1211 |
| balanced divide | MRE | balanced | ✓ | × | 9.63 | −0.1991 | 4.21 | −0.1403 | 2.93 | −0.0312 | 3.15 | −0.1781 | 3.62 | −0.2655 | 2.97 | −0.1620 |
|  |  |  |  | default | 6.15 | −0.0017 | 3.63 | −0.0073 | 2.94 | +0.0018 | 2.80 | −0.0053 | 3.09 | −0.0110 | 2.92 | +0.0218 |
|  |  |  |  | extended | **5.61** | +0.0034 | 3.83 | +0.0056 | 3.36 | +0.0045 | 3.18 | +0.0010 | 3.61 | +0.0055 | 3.98 | **+0.1266** |
| no bipartition test | MRE | quartet | × | × | 7.12 | −0.0454 | 3.87 | −0.0169 | 3.05 | −0.0221 | 2.91 | −0.0173 | 3.04 | −0.0292 | 3.13 | −0.0927 |
|  |  |  |  | default | 5.92 | −0.0009 | 3.66 | +0.0016 | 2.99 | +0.0000 | = | +0.0000 | = | +0.0000 | = | +0.0000 |
|  |  |  |  | extended | 5.68 | +0.0022 | 3.82 | +0.0067 | 3.38 | +0.0036 | 3.21 | +0.0010 | 3.60 | +0.0072 | 4.08 | +0.1252 |
| true tree |  |  |  |  |  | −0.0086 |  | −0.0432 |  | −0.0462 |  | −0.0189 |  | −0.1084 |  | +0.0745 |
| ASTRAL-X |  | S1 |  |  | 54.01 | −4.0960 | 10.04 | −1.0542 | 4.09 | −0.0783 | 9.65 | −1.9196 | 12.79 | −3.3699 | 7.60 | −2.4608 |
|  |  | S2 |  |  | 7.51 | −0.1643 | 4.30 | −0.0525 | 3.46 | +0.0040 | 3.75 | −0.1454 | 4.03 | −0.2594 | 4.66 | −0.1275 |
|  |  | S3 |  |  | 5.89 | −0.0017 | 3.91 | +0.0073 | 3.46 | +0.0048 | 3.32 | +0.0012 | 3.72 | +0.0049 | 4.48 | +0.0691 |
| STELAR-X |  | S1 |  |  | 56.72 | −4.5262 | 11.70 | −1.4487 | 4.42 | −0.1917 | 9.97 | −2.3531 | 13.27 | −3.9277 | 7.86 | −3.1897 |
|  |  | S2 |  |  | 11.07 | −0.4969 | 5.45 | −0.2156 | 3.80 | −0.0532 | 4.13 | −0.2784 | 5.07 | −0.5651 | 5.62 | −0.4835 |
|  |  | S3 |  |  | 8.07 | −0.2443 | 4.48 | −0.0699 | 3.79 | −0.0609 | 3.78 | −0.0822 | 3.96 | −0.1296 | 5.64 | −0.4574 |
| ASTRAL-MP |  |  |  |  | 5.87 | −0.0001 | 3.90 | +0.0064 | 3.50 | +0.0048 | 3.37 | +0.0010 | 3.72 | −0.0029 | 4.54 | +0.0800 |
| ASTRAL-IV |  |  |  |  | 5.74 | **+0.0052** | 3.88 | **+0.0083** | 3.54 | **+0.0063** | 3.36 | **+0.0016** | 3.79 | **+0.0156** | n.r. | n.r. |
| wQFM-TREE |  |  |  |  | 5.79 | −0.0190 | 3.61 | −0.0078 | 2.98 | −0.0412 | 2.80 | −0.0325 | 3.31 | −0.0576 | n.r. | n.r. |

#### Starting tree

The MRE tree alone was accurate at low ILS and at 10,000 taxa (where ILS is presumably low as well), where its error was below QRAFT’s. But it failed under high ILS with 2.5× QRAFT’s error, and still 1.8 ×even with extended (NNI and SPR) refinement. The full pipeline is therefore much more robust to ILS than the MRE consensus. Still, the MRE tree’s reasonable accuracy shows that it is a sound place to start. Its quartet score, however, was far below QRAFT’s, by up to 34% of the gene-tree quartets at 10,000 taxa. This gap mostly reflects its polytomies, which resolve no quartets. For the same reason, a missing bipartition counts half as much as a wrong one in its RF rate. The RF rate therefore penalizes polytomies less than the quartet score does. Where a binary tree is required, however, the MRE tree alone cannot provide one, and QRAFT’s divide and regrafting can also be seen as a cheap way to resolve an unresolved consensus or guide tree with the quartet objective.

Starting from the refined MRE tree changed QRAFT’s default RF rate by at most 0.16% on ASTRAL-II and raised it from 3.12% to 3.55% at 10,000 taxa. A better starting tree thus brings little, which suggests that the divide needs only a few good MRE edges supported by the gene trees, not a fully correct starting tree.

#### Divide and bipartition test. The balanced divide, which skips the quartet score of candidate edges, was competitive at low ILS and at 10,000 taxa. Without refinement, however, it raised the RF rate from 6.92% to 9.63% under high ILS and from 3.81% to 4.21% under medium ILS, and refinement narrowed but did not close this gap (6.15% against 5.77% under high ILS). Choosing the edge with the highest score *s*(*A B*) therefore matters most when the MRE tree is least reliable, since the edge QRAFT tests is then the one the gene-tree quartets support best. The bipartition test helped the default pipeline and never hurt by more than 0.02%: removing it raised the RF rate under high ILS from 6.92% to 7.12% without refinement and from 5.77% to 5.92% with default refinement. With 1,000 genes, fewer than *K*_0_, the test was permissive (*t_xy_* < 0, Sect. 4). It becomes stricter with more genes, so its effect may be larger there

#### Other methods

A QRAFT variant had the lowest RF rate in every column, and QRAFT’s default tree was more accurate than every other method, except ASTRAL-IV under high ILS (5.74% against 5.77%). ASTRAL-IV, however, reached a slightly higher quartet score than QRAFT’s default on all ASTRAL-II columns, by 0.002 to 0.016% of the gene-tree quartets, a small shortfall of QRAFT’s search that we acknowledge. As the true tree’s lower score on ASTRAL-II shows, a higher quartet score did not mean a lower RF rate here.

## S3 Proof of Statistical Consistency

### S3.1 Setting and proof overview

Let *X* be a fixed set of *n* ≥ 4 taxa and let G = {*g*_1_, …, *g_k_*} be the observed unrooted gene trees, where *g_i_* has taxon set *X_i_* ⊆ *X*. We use the notation of the problem definition in the main paper: *g_i_* is the observed tree, denoted by *H_g_* in the description of the bipartition test. Write 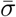 for the unrooted topology of the species tree *σ*. All limits below are taken as *k* → ∞, with *X, σ* and the taxon-presence distribution fixed.

#### Definition S3.1

(Model conditions). We assume the model conditions of Definition 2.4:

(M1) *Multispecies coalescent:* the unrestricted gene trees are independent and identically distributed under the MSC on a binary species tree *σ* with positive internal branch lengths. The observed tree *g_i_* is the restriction of the corresponding gene tree to *X_i_*.

(M2) *Full subset coverage:* the sets *X_i_* are independent and identically distributed, independently of the unrestricted gene trees, and Pr(*B* ⊆ *X_i_*) > 0 for every *B* ⊆ *X* with |*B*| ≤ 4.

Condition (M2) is the full subset coverage model of Nute et al. [25] for subsets of up to four taxa. Although QRAFT also accepts unresolved input trees, the consistency proof concerns the observed restrictions of MSC gene trees, which are binary whenever they contain at least three taxa. Under these conditions, we prove Theorem 2.5: QRAFT returns 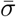 with probability tending to one, for every starting tree, with or without NNI and SPR refinement.

#### Proof overview

QRAFT constructs a tree by accepting candidate bipartitions, recursively reconstructing their two sides, and joining the resulting subtrees through contextual regrafting. It may then apply NNI and SPR refinement. We show that, with probability tending to one, the bipartition test accepts exactly the true bipartitions (Lemma S3.6), contextual regrafting joins correctly reconstructed subtrees at their true attachment edges (Lemma S3.8), and refinement cannot move away from the true species tree (Corollary S3.8.1). Induction over the recursive subproblems then establishes Theorem 2.5.

The main challenge is the bipartition test. Nute et al. [25] established the consistency of exact ASTRAL under (M1) and (M2), using the fact that deleting taxa outside a quartet does not change its topology. In contrast, the inter-anchor subpath lengths used by our test can change when taxa are deleted, as can the internode distances used by NJst and ASTRID [26, 27]. We first show that the expected gap nevertheless characterizes the species-tree quartet. We then establish concentration around this expectation and show that an appropriate threshold separates true and false bipartitions.

### S3.2 Expected gap and concentration

#### Inter-anchor subpath gap

Let *A*|*B* be a candidate bipartition of a subproblem *S* ⊆ *X*, with |*A*|, |*B*| ≥ 2, and fix anchors *a* ∈ *A* and *b* ∈ *B*. Let 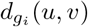 be the number of edges between *u* and *v* in the whole observed gene tree *g_i_*, after suppressing degree-two vertices. As in Sect. 2.4, for a gene containing *a, b, c*, define

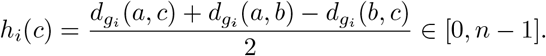

This is the position at which *c* attaches to the *a*–*b* path, measured from *a*. For distinct non-anchor taxa *x, y*, define

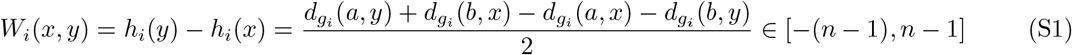

when {*a, b, x, y*} ⊆ *X_i_*, and set *W_i_*(*x, y*) = 0 otherwise. Thus *W_i_*(*x, y*) is positive when *y* attaches later than *x* along the path from *a* to *b*, negative when it attaches earlier, and zero when they attach at the same vertex. Distances are computed on the whole observed genes, including taxa outside *S*. Consequently, a comparison depends only on (*a, b, x, y*), not on the subproblem containing these taxa.

Let *Z_i_* = 1[{*a, b, x, y*} ⊆ *X_i_*] and define

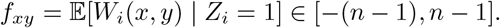

The notation suppresses the anchors *a, b*. The expectation is well defined by (M2) and is the same for every gene *i*.

##### Lemma S3.2

(Sign of the expected gap). *There is a constant f*_*_ > 0 *such that, for every four distinct taxa a, b, x, y, f_xy_* ≥ *f*_*_ *if* 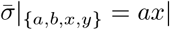*by, f_xy_* = 0 *if it is ab*|*xy, and f_xy_* ≤ −*f*_*_ *if it is ay*|*bx.*

*Proof.* Fix *Y* ⊇ {*a, b, x, y*} with Pr(*X_i_* = *Y*) > 0. By (M2), conditioning on *X_i_* = *Y* does not change the distribution of the unrestricted gene tree. Sampling consistency of the MSC therefore gives *g_i_* the MSC distribution on *σ*|*_Y_*. By Theorem 4.2 of Allman, Degnan and Rhodes [28], the expected edge-count distances

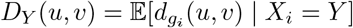

form an additive distance matrix for 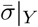, with strictly positive internal edge weights. These weights need not equal the species-tree branch lengths in coalescent units.

Restrict this expected-distance tree to the four taxa *q* = {*a, b, x, y*}, and let 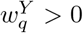 be its internal edge length. By (S1) and the four-point identity for additive distances,

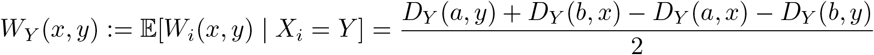

Satisfies

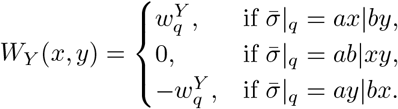

The quartet topology is the same for every such *Y*. By the law of total expectation,

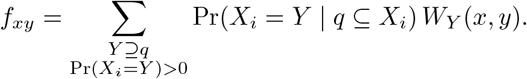

The coefficients are nonnegative and sum to one. Hence, defining

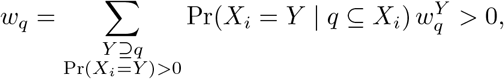

we obtain *f_xy_* = *w_q_*, 0 or −*w_q_* for the three respective quartet topologies. Finally,

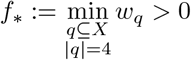

is a minimum over finitely many positive numbers, proving the uniform bounds.□

**Shared genes.** To estimate *f_xy_*, we use the shared genes

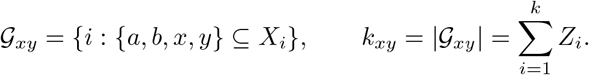

By (M2),

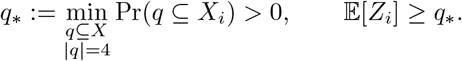

##### Proposition S3.3

(Shared genes). *With probability tending to one, k_xy_* ≥ *q*_*_*k/*2 > 0 *for every choice of distinct anchors a, b and distinct non-anchor taxa x, y. More precisely,*

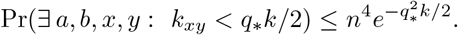

*Proof.* The indicators *Z_i_* are independent by (M2), and E[*k_xy_*] ≥ *q*_*_*k*. Hoeffding’s inequality gives

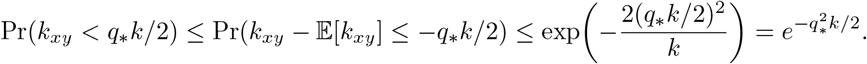

A union bound over the at most *n*^4^ ordered choices of four distinct taxa proves the claim.□

#### Mean gap

For *k_xy_* > 0, the empirical mean gap is

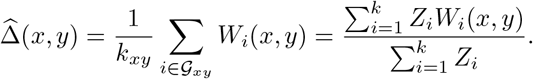

For the probability statements below, set 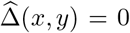 when *k_xy_* = 0. This convention does not affect the limit, by Proposition S3.3. Put *R* = 2(*n* − 1), the width of the interval in (S1).

##### Lemma S3.4

(Concentration of the mean gap). *For every* 0 <*ε* <*R,*

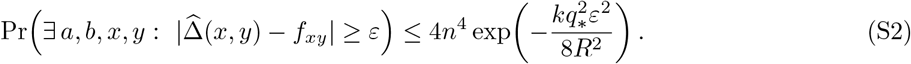

*In particular,* 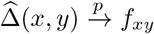 *for every comparison.*

*Proof.* Fix *a, b, x, y*. The pairs (*X_i_, g_i_*) are independent and identically distributed by (M1) and (M2). Moreover,

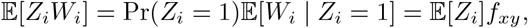

so E[*Z_i_*(*W_i_* −*f_xy_*)] = 0. The variable *Z_i_*(*W*_*i* −_ *f_xy_*) lies in [− (*n* − 1) −*f_xy_, n* − 1 −*f_xy_*], an interval of width *R* containing zero. Hoeffding’s inequality therefore gives

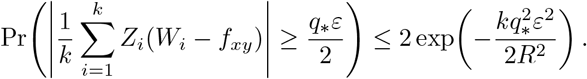

Applying Hoeffding’s inequality to ∑*_i_ Z_i_*, as in Proposition S3.3, also gives

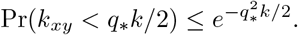

If neither event occurs, then

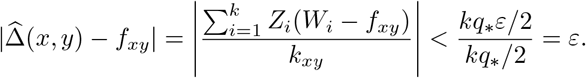

Thus, by the union bound and *ε* <*R*,

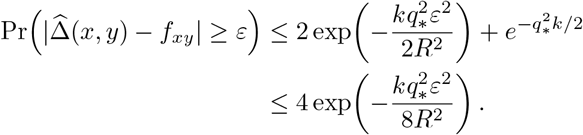

A union bound over the at most *n*^4^ ordered choices of four distinct taxa proves (S2). Its right-hand side tends to zero for every fixed 0 <*ε* <*R*.

*Proof of Lemma 2.1.* Lemma S3.4 gives 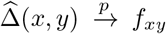, and Lemma S3.2 gives the three possible signs and the uniform positive margin *f*_*_. Together these prove the mean gap sign characterization in the main paper. □

### S3.3 Consistency of the bipartition test

#### The test

For fixed anchors *a* ∈ *A* and *b* ∈ *B*, the desired extreme taxa are *x*^*^ ∈ *A* \ {*a*} attaching last on the species-tree *a*–*b* path and *y*^*^ ∈ *B* \ {*b*} attaching first (Fig. 2). QRAFT searches for these taxa by scanning each side. On *A* \ {*a*}, it starts with a non-anchor taxon *c* and replaces *c* with *x* whenever 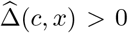. On *B b*, it replaces *c* with *y* whenever 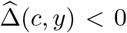. An empirical tie retains the current taxon. We show below that these searches return taxa at the desired extreme positions with probability tending to one. For brevity, write *x*^*^, *y*^*^ for the returned taxa as well. The test accepts *A*|*B* when

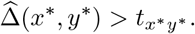

The threshold must tend to zero more slowly than the sampling noise. We formalize the two properties used in the proof.

##### Definition S3.5

(Admissible threshold). For fixed *X*, a threshold rule is *admissible* if, as *k_xy_* → ∞,

(T1) *t_xy_* → 0, and

(T2) 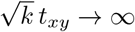, uniformly over all comparisons and input configurations. Explicitly, for every *η* > 0 and *M* > 0, there is an integer *j*_0_ such that 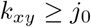 implies |*t_xy_*| <*η* and 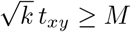 for every such configuration. The threshold may depend on the observed presence sets. In particular, (T2) requires it to be positive for sufficiently large shared counts.

##### Lemma S3.6

(Consistency of the bipartition test). *Under the model conditions and an admissible threshold, the bipartition test accepts exactly the true bipartitions for every subproblem, candidate and pair of anchors with probability tending to one.*

*Proof.* The test succeeds if (i) every comparison has shared genes, (ii) the two searches return taxa at the extreme species-tree attachment positions, and (iii) the gap at the extreme pair exceeds the threshold exactly for true bipartitions.

Let *H_k_* be the event that *k_xy_* ≥ *q*_*_*k/*2 for every comparison. By Proposition S3.3,

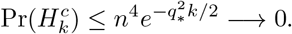

Let *E_k_* be the event that *k_xy_* > 0 and 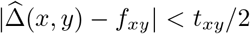 for every comparison. Since *t_xy_* may depend on the presence sets, we do not substitute it directly for *ε* in (S2). Instead, fix *M* > 0. On *H_k_*, admissibility implies that, for all sufficiently large *k*, every comparison satisfies

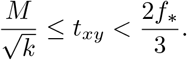

For such *k*, also taking 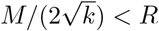, Lemma S3.4 gives

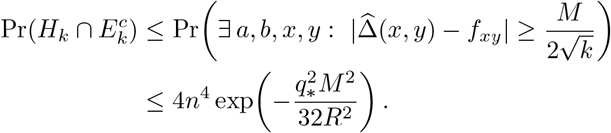

Consequently,

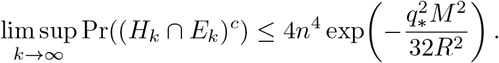

This holds for every *M* > 0. Letting *M* → ∞ proves Pr(*H_k_* ∩ *E_k_*) → 1.

Assume now that *H_k_* ∩ *E_k_* holds and *k* is sufficiently large that 0 <*t_xy_* < 2*f*_*_*/*3 for every comparison. Then *k_xy_* ≥ *q*_*_*k/*2 > 0, proving (i), and

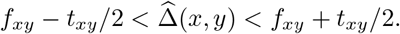

*(ii) The searches find the extreme positions.* By Lemma S3.2, two taxa at distinct species-tree attachment positions have an expected gap of magnitude at least *f*_*_. Hence

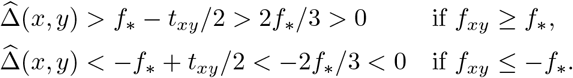

Thus every comparison between distinct attachment positions has the correct sign. During the scan of *A\*{*a*}, the current taxon is at the latest position among those scanned so far: a taxon at a later position replaces it, and a taxon at an earlier position cannot replace it. A taxon at the same position may replace it without changing the position. This proves the invariant by induction over the scan. The symmetric argument for *B\*{*b*} gives its earliest position. No sign condition is needed when *f_xy_* = 0. Therefore, the searches return valid extreme taxa *x*^*^, *y*^*^, proving (ii).

*(iii) The extreme pair distinguishes true and false bipartitions.* If *A*|*B* is a true bipartition of 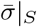, its separating edge lies between every *A* attachment and every *B* attachment. Every cross pair *x* ∈ *A* \ {*a*}, *y* ∈ *B* \ {*b*} therefore displays *ax*|*by* in the species tree, so

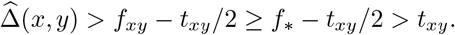

In particular, the extreme pair passes.

Conversely, if the last *A* attachment strictly preceded the first *B* attachment, an edge of the anchor path between them would separate all taxa of *A* from all taxa of *B*, realizing *A*|*B* in 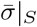 Thus, for a false bipartition, *y*^*^ attaches at or before *x*^*^. If their positions coincide, they lie in the same pendant subtree of the binary species tree and induce *ab*|*x*^*^*y*^*^; if *y*^*^ attaches earlier, they induce *ay*^*^|*bx*^*^. Lemma S3.2 gives *f_x_*\**_y_^*^* ≤ 0 in either case, and hence

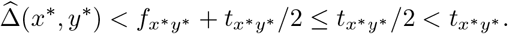

The false candidate fails. A singleton side is accepted directly and always defines a true bipartition.

The events above cover all ordered choices of *a, b, x, y* in *X*. Because the comparisons use whole observed genes, the same events apply simultaneously to every subproblem and every adaptively chosen candidate or anchor pair. This proves the lemma.□

### S3.4 Regrafting, refinement and the consistency theorem

The bipartition test ensures that division uses true bipartitions. It remains to show that the quartet score joins the reconstructed sides correctly and preserves the species tree under refinement. For a tree *T* on *S* ⊆ *X*, write

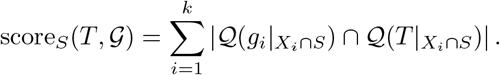

Restrictions to fewer than four taxa contribute zero. This agrees with the main paper’s quartet score when *S* = *X*.

#### Lemma S3.7

(Species quartets dominate). *With probability tending to one, 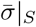 uniquely maximizes* score*_S_*(*T*, G) *among binary trees on S, simultaneously for every nonempty S* ⊆ *X.*

*Proof.* Fix a four-taxon set *q* ⊆ *X*, and let *τ_q_* > 0 be its induced internal branch length in coalescent units. By (M2) and sampling consistency of the MSC, conditional on *q* ⊆ *X_i_*, the quartet *g_i_*|*_q_* displays the species resolution with probability 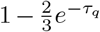 and each alternative with probability ^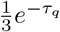^ [29]. For either alternative resolution 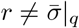, define

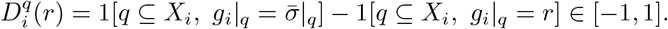

This variable is 1 for an observed species quartet, −1 for the alternative *r*, and 0 if *q* is absent or the other alternative is displayed. The variables are independent and identically distributed across genes, with

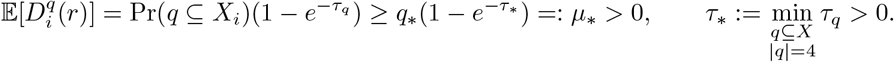

Hoeffding’s inequality, using the range width 2, gives

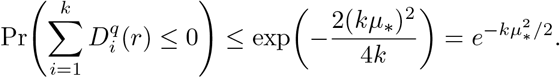

Let *Q_k_* be the event that 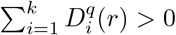 for every four-taxon set *q* and each alternative 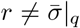. A union bound gives

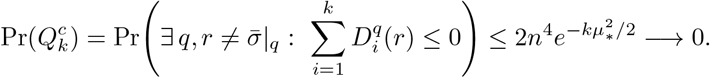

Assume *Q_k_* holds. For any *S* ⊆ *X* and any binary tree *T* on *S*, the quartet score is

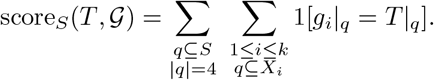

If |*S*|≥ 4 and 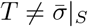, then *T* differs from 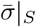 on at least one quartet, because a binary tree is determined by its quartets. Therefore,

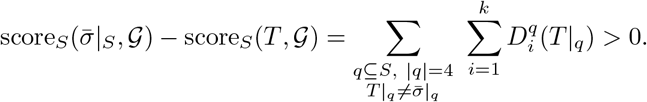

Every summand indexed by a differing quartet is positive on *Q_k_*, and there is at least one. Thus 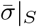 is the unique maximizer for every *S*. For 1≤ |*S*| ≤3, the unrooted topology is unique. This is the quartet argument underlying the consistency of exact ASTRAL [30, 31], extended to missing taxa as in [25].

#### Lemma S3.8

(Contextual regrafting). *Let A*|*B be a true bipartition of a subproblem S, with correctly reconstructed subtrees* 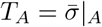 *and* 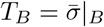*. On Q_k_, contextual regrafting returns* 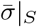.

*Proof.* Joining *T_A_* and *T_B_* at their species-tree attachment edges yields 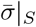. By Lemma 2.2, the quartet score of a joined tree is *c* + *ϕ_A_*(*e_A_*) + *ϕ_B_*(*e_B_*), so contextual regrafting maximizes it exactly by optimizing the two attachment choices independently. When one side is a singleton, only the attachment edge on the other side must be chosen. In every case, 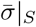 is among the possible joins and, on *Q_k_*, it is the unique quartet-score maximizer by Lemma S3.7. Therefore, contextual regrafting returns it.

#### Corollary S3.8.1

(Refinement keeps the species tree). *On Q_k_, NNI and SPR refinement started from* 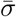 *return* 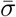.

*Proof.* By Lemma S3.7, every topology different from 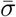 has a strictly smaller quartet score. Refinement applies only moves with positive exact quartet-score gain. Hence no NNI or SPR move can be applied from 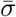. Screening rules, support gates and limits on the number of moves cannot change this conclusion.

*Proof of Theorem 2.5.* Let B*_k_* be the event that the bipartition test accepts exactly the true bipartitions for every subproblem, candidate and anchor pair. Lemmas S3.6 and S3.7 give Pr(ℬ*_k_*) →1 and Pr(*Q_k_*) → 1 for any admissible threshold.

Assume ℬ *_k_* ∩ *Q_k_* holds. For any starting tree, we show by induction on |*S*| that the recursive call on *S* returns 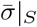. The claim is immediate for |*S*| ≤ 3, since the topology is unique. For |*S*| ≥ 4, every accepted candidate *A*|*B* is a bipartition of *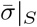* on ℬ*_k_*. If no candidate is accepted or available, QRAFT uses a trivial bipartition separating one taxon from the rest, which also belongs to 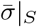. Since both sides are nonempty and strictly smaller than *S*, the induction hypothesis gives 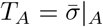 and 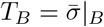. Lemma S3.8 then gives 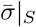 after contextual regrafting.

At the root, *S* = *X*, so QRAFT returns 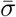 before refinement. Corollary S3.8.1 shows that NNI and SPR refinement preserve it. Writing *T_k_* for the final output,

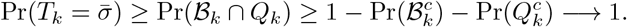

The result is independent of the starting tree. The next subsection verifies that QRAFT’s implemented threshold is admissible and that its additional comparison and anchor-selection rules preserve the argument. □

### S3.5 Implementation details

#### QRAFT’s threshold

For fixed constants *C* > 0 and *K*_0_ ≥ 0, QRAFT uses

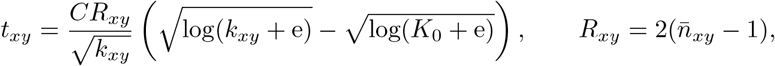

where 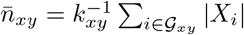 is the mean number of taxa in the whole shared genes, including taxa outside the subproblem. The threshold is negative for *k_xy_* <*K*_0_, zero at *K*_0_, and positive above *K*_0_. Its permissive behavior at small shared counts does not affect admissibility.

##### Lemma S3.9

(QRAFT’s threshold is admissible). *QRAFT’s threshold satisfies* (*T1*)*and* (*T2*)*of Definition S3.5.*

*Proof.* Every shared gene contains at least four taxa and at most *n*, so 6 ≤ *R_xy_* ≤ *R* = 2(*n* − 1). For *j* = *k_xy_* >*K*_0_,

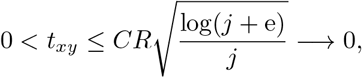

proving (T1). Also, since *k* ≥ *j* and the threshold is positive,

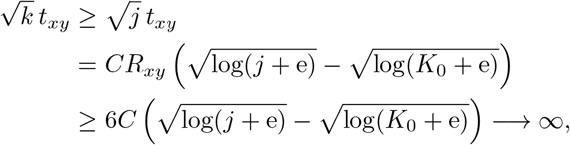

proving (T2). Both bounds depend only on *j* and fixed constants, so the convergence is uniform over all comparisons and input configurations. □

**Additional comparisons.** For the chosen anchors, QRAFT tests the pairs

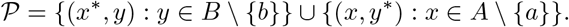

It tests (*x*^*^, *y*^*^) first, followed by the remaining pairs in the two sets, counting the extreme pair only once. Pairs with no shared gene are skipped. The anchor pair accepts the candidate exactly when the smallest tested margin 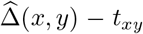 is strictly positive. On *H_k_* ∩*E_k_* for sufficiently large *k*, every cross pair of a true bipartition passes by part (iii) of the proof of Lemma S3.6, whereas a false bipartition already fails at (*x*^*^, *y*^*^). Every comparison is defined on *H_k_*, so none is skipped there.

#### Anchor selection and unavailable tests

The anchor candidates are the two taxa on each side present in the most genes, with ties broken by their order in the active taxon set. For each anchor pair, the searches start with the first non-anchor taxon on each side present in a gene with both anchors. An empirical tie or a comparison with no shared gene retains the current taxon. An anchor pair is unusable if either side has no such non-anchor taxon, or if the two taxa returned by the searches share no gene with both anchors. QRAFT accepts the candidate if any tested anchor pair accepts it. If no anchor pair is usable, the test is unavailable and the candidate is accepted. This permissive finite-sample rule cannot occur on *H_k_*, where every four-taxon set has shared genes and every anchor pair for a nontrivial candidate is usable. Since Lemma S3.6 holds simultaneously for all anchor choices, selecting anchors from the data does not affect consistency. A singleton side is accepted directly. If no candidate is available or the configured candidate proposals are exhausted without acceptance, QRAFT uses a trivial bipartition, as required by the induction above.

## Notes

https://github.com/risenfromashes/qraft

### Competing Interest Statement

The authors have declared no competing interest.

